# Common Ancestry of the *Id* Locus: Chromosomal Rearrangement and Polygenic Possibilities

**DOI:** 10.1101/2023.12.29.573606

**Authors:** Ashutosh Sharma, Nagarjun Vijay

## Abstract

The diversity in dermal pigmentation and plumage color among domestic chickens is striking, with Black Bone Chickens (BBC) particularly notable for their intense melanin hyperpigmentation. This unique trait is driven by a complex chromosomal rearrangement on chromosome 20 at the *Fm* locus, resulting in the overexpression of the *EDN3* (a gene central to melanocyte regulation). In contrast, the inhibition of dermal pigmentation is regulated by the *Id* locus. Although prior studies using genetic crosses, GWAS, and gene expression analysis have investigated the genetic underpinnings of the *Id* locus, its precise location and functional details remain elusive. Our study aims to precisely locate the *Id* locus, identify associated chromosomal rearrangements and candidate genes influencing dermal pigmentation, and examine the ancestral status of the *Id* locus in BBC breeds. Using public genomic data from BBC and non-BBC breeds, we refined the *Id* locus to a ∼1.6 Mb region that co-localizes with Z amplicon repeat units at the distal end of the q-arm of chromosome Z within a 10.36 Mb inversion in Silkie BBC. Phylogenetic and population structure analyses reveal that the *Id* locus shares a common ancestry across all BBC breeds, much like the *Fm* locus. Selection signatures and highly differentiated BBC-specific SNPs within the *MTAP* gene position it as the prime candidate for the *Id* locus with *CCDC112* and additional genes, suggesting a possible polygenic nature. Our results suggest that the *Id* locus is shared among BBC breeds and may function as a supergene cluster in shank and dermal pigmentation variation.

## Introduction

Phenotypic diversity in pigmentation patterns within avian species has long been a focal point of biological research. The chicken has garnered considerable attention due to its economic and culinary significance and the remarkable variety of plumage colors and dermal pigmentation exhibited across different breeds. The most striking of such pigmentation phenotypes, fibromelanosis (*Fm*), refers to hyperpigmentation throughout the tissues and organs, encompassing the dermal layer of skin, bones, muscles, nerves, and gonads of black-bone chicken (BBC) (Dorshorst et al. 2011; Shinomiya et al. 2012). The underlying genetic cause of this hyperpigmentation, often referred to as the "black bone phenotype," was first investigated by Bateson and Punnett using the Silkie BBC breed (Bateson et al. 1911). Genetic crossing experiments identified two independent epistatically interacting factors influencing the degree of pigmentation observed in hybrid crosses between Silkie and White Leghorn chickens (Dunn and Jull 1927; Arora et al. 2011; Tian et al. 2014a). These two genetic factors responsible for dermal pigmentation are termed "fibromelanosis (*Fm*) locus" and "inhibitor of dermal melanin (*Id*) locus" (Mukherjee et al. 1969; Tixier-Boichard 2002).

The genetic basis of the *Fm* locus has been identified on chromosome 20 as a complex chromosomal rearrangement encompassing three different non-paralogous genomic regions, Dup1, Int, and Dup2, with the Dup1 and Dup2 regions being duplicated (Dorshorst et al. 2011; Shinomiya et al. 2012; Dharmayanthi et al. 2017; Sohn et al. 2018; Shinde et al. 2023). The most important region Dup1 contains five protein-coding genes: *EDN3*, *ZNF831*, *SLMO2*, *ATP5E*, and *TUBB1*. The prominent factor contributing to the intense melanization observed in the internal organs of BBC is the heightened expression of the Endothelin 3 gene (*EDN3*), which plays a pivotal role in producing melanocytes, specialized cells responsible for melanin synthesis (Baynash et al. 1994). The increased expression of *EDN3* resulting from the rearrangement is a key determinant in the hypermelanization phenomenon (Dorshorst et al. 2011).

BBC breeds are found globally, with a particularly strong presence in Asia. Comprehensive research, including genetic diversity studies, genome-wide analyses, and mitochondrial DNA evidence, indicates that China is the primary origin of all BBC breeds (Zhang et al. 2018; Huang et al. 2021). Interestingly, BBC breeds from diverse geographic regions across Asia share a unique chromosomal rearrangement at the *Fm* locus (Shinde et al. 2023). A chromosome-level denovo genome assembly of a Silkie BBC individual proposed an alternative rearrangement (*\*Fm_1*) at the *Fm* locus (Zhu et al. 2023). However, a recent reanalysis of this data and additional data from other BBC breeds supports the presence of the common rearrangement (*\*Fm_2*) at the *Fm* locus (Sharma and Vijay 2024). The shared rearrangement and the population genetic signature of this region suggest a common ancestral origin for all BBC breeds.

In contrast to the autosomal dominant *Fm* locus, the mode of inheritance of *Id* is sex-linked incomplete dominance. As its name suggests, the *Id* locus is known to inhibit dermal pigmentation to different degrees based on its genetic state. Gene mapping studies have identified the physical location of the *Id* locus at the distal end of the q-arm of chromosome Z, in proximity to the sex-linked barring loci B (Bitgood 1985, 1988; Dorshorst and Ashwell 2009). The availability of genome assemblies helped identify that the chicken Z chromosome has a distinctive genetic structure known as the Z amplicon repeat unit (ZARU) at the distal end of the q-arm. The Z amplicon repeats consist of multiple copies of four genes: *ADCY10*, *C2orf3*, *ARHGAP33*, and *MRPL19* (Bellott et al. 2010; Yan et al. 2010) interspersed with tandem repeats (Edeen 1985).

Early draft genomes supported the localization of the *Id* locus to the distal end of the q-arm of the Z chromosome (Dorshorst et al. 2010). As genomic resources improved, the position of the *Id* locus was further refined. In recent years, utilizing a genome-wide SNP-trait association analysis, researchers pinpointed a potential *Id* locus in the galGal4 genome assembly, situated approximately at 78.4 Mb on chromosome Z (Dorshorst et al. 2010; Siwek et al. 2013; Li et al. 2014). Several studies have been undertaken involving crosses between BBC and non-BBC breeds, all in pursuit of identifying the candidate *Id* gene. These investigations have reported various potential genes which are associated with dermal and shank pigmentation variations (Mcgibbon 1978; Arora et al. 2011; Li et al. 2014; Tian et al. 2014b; Xu et al. 2017; Cha et al. 2023).

Numerous studies have been conducted to unravel the mysteries of the *Id* locus. Yet, the puzzle surrounding this locus persists, and the precise genomic region or gene responsible for its function remains elusive. However, with recent improvements in chicken genome assemblies and access to population genomic data from various BBC and non-BBC breeds, there is now an opportunity for a more comprehensive analysis to identify the *Id* locus accurately. Our study aims to address the following objectives: **1)** Assess the co-localization of Z amplicon repeats and *Id* locus along the Z chromosome. **2)** Evaluate the ancestral relationship at the *Id* locus among different BBC breeds. **3)** Identification of structural variation between BBC and non-BBC. **4)** Identify BBC-specific genomic signatures within the *Id* locus using population genomic statistics. **5)** Prioritize genomic variants associated with dermal pigmentation.

These objectives will guide a detailed investigation to advance our understanding of the *Id* locus and its role in pigmentation.

## Materials and methods

### Data collection

To comprehensively investigate the genomic structure and diversity of the *Id* locus, we acquired publicly accessible whole-genome resequencing short-read Illumina data of 270 chicken individuals. This dataset comprises three distinct wild junglefowl species: green, Ceylon, and grey. In addition to the wild junglefowl, the dataset includes multiple breeds of domesticated chicken. We included various BBC breeds, including Kadaknath, Silkie, Yeonsan Ogye, and Chinese breeds. Additionally, we incorporated non-black native chicken breeds like Aseel, Japanese gamefowl, Chinese gamefowl, Tibetan native chicken, Ethiopian native chicken, Sri Lankan native chicken, and Korean native chicken. The collection also encompasses commercial chicken breeds like White leghorn, Broiler, and Rhode Island Red.

Of the 270 individuals used in this study, 42 belonged to the BBC category, while 228 were categorized as non-BBC. We sourced these data from public repositories such as the National Center for Biotechnology Information (NCBI), the European Nucleotide Archive (ENA), the National Genomics Data Center (NGDC), and the Korean National Agricultural Biotechnology Information Center (KNABIC). The genome coverage of the resequencing data for each individual was at least more than 10x. Comprehensive details for each individual can be found in Table S1.

### Evaluation of assembly quality at the distal end of chromosome Z

To ensure the high quality of genome assemblies at the distal end of the q-arm of chromosome Z, we conducted assembly validation in the region of interest, specifically focusing on the potential *Id* locus. For this, we downloaded the high-coverage chicken ONT long-read data (SRR15421342, SRR15421343, SRR15421344, SRR15421345, SRR15421346, SRR13494713, and SRR13494714) of Huxu and Arbor Acres breeder rooster breed from ENA. We mapped these data to Huxu (GCA_024206055.2), GRCg6a (GCA_000002315.5) and GRCg7b (GCA_016699485.1) genome assemblies using the minimap2 v2.27-r1193 (Li, 2018) with -- secondary=no --sam-hit-only --MD -Y -t 24 -ax map-ont flags.

To validate the continuity and integrity of these assemblies, we visually inspected a tiling path of long reads spanning the *Id* locus region using the UCSC genome browser (Karolchik et al. 2003; Fujita et al. 2011) for all three genome assemblies. To further evaluate the integrity of these genome assemblies, we used the Klumpy tool (Madrigal et al. 2024) on the aligned long reads using the “scan alignment” flag. We also compared all three genome assemblies, using SyRI (Synteny and Rearrangement Identifier) (Goel et al. 2019) to explore the structural similarities and rearrangements. To visualize the results obtained from SyRI, we utilized plotsr (plot structural rearrangements) (Goel and Schneeberger 2022). Our analysis found that the GRCg6a has better assembly quality (see **results** for details) compared to GRCg7b in the region of interest (*Id* locus). Therefore, we used GRCg6a as the reference genome assembly for subsequent studies.

### Read mapping and variant calling

We mapped the paired-end raw Illumina reads from 270 individuals to the Gallus_gallus.GRCg6a genome assembly. The mapping was executed using the "bwa mem" flag, employing the default parameters within the BWA v 0.7.17-r1188 (Li and Durbin 2009). We incorporated read group information to enhance data integrity and eliminated duplicate reads from the resulting BAM files, utilizing the Picard tools (accessible at https://github.com/broadinstitute/picard). We employed two distinct tools for variant calling: bcftools version 1.10.2 (Li and Durbin 2009; Danecek et al. 2021) and FreeBayes version 1.0.2 (Garrison and Marth 2012). For bcftools, we used various quality control flags such as -mapping quality -C 50, -min base quality -Q 20, and -min mapping quality -q 20. Similarly, for the FreeBayes, we imposed various quality control flags, including -min-alternate-count -C 10, -min-mapping-quality -m 20, -min-base-quality -q 20, and -min-coverage, requiring a minimum of 10 reads at a given locus. The aim was to identify SNPs with high quality and robustness. To refine the dataset, we applied stringent filters using VCFtools version 0.1.16 (Danecek et al. 2011) to exclude low-quality variant calls and indels using the flags --remove-indels, --max-alleles 2, and --max-missing 0.95. Additionally, to enhance our SNP dataset’s precision, we extracted common SNPs concordantly identified by both variant callers, minimizing the possibility of false positive SNP calls. After the filtration and obtaining the common SNPs, we ended with 62,102,065 SNPs genome-wide and 3,397,676 for chromosome Z.

### Annotation of ZARU at the distal end of the q-arm of chromosome Z

In light of previous research (Bellott et al. 2010) about Z amplicon repeat units, we employed two distinct methodologies: (1) blastn and (2) read coverage estimation, to annotate the Z amplicon repeat units (ZARU) at the distal end of q-arm of chromosome Z in the GRCg6a genome assembly.

In the first approach, we utilized the blastn v2.9.0 (Altschul et al. 1990) to elucidate the spatial distribution of four specific genes: *ADCY10*, *C2orf3*, *ARHGAP33*, and *MRPL19*. To achieve this, we used the genomic sequences of all the annotated isoforms of these genes in the latest chicken genome as query sequences from NCBI for performing BLAST searches against the GRCg6a genome assembly. Subsequently, the obtained BLAST hits were saved in bed format, and overlapping hits were merged using the merge command in bedtools v2.27.1(Quinlan and Hall 2010). To visualize the distribution pattern of these genes, we divided chromosome Z into 1 Kb non-overlapping windows using bedtools. For each of these windows, we calculated the coverage of the BLAST hits using the bedtools coverage command.

In our second approach, we took a different perspective by evaluating read coverage in 10 Kb non-overlapping windows spanning chromosome Z. This allowed us to analyze read counts within each window, providing valuable confirmation regarding the presence of multiple copies of the four genes. Both methods were designed to yield robust evidence for annotating the ZARU regions. We visualized the distribution pattern and read coverage of ZARU regions using R v4.4.1 (Ihaka and Gentleman 1996), which allowed us to confirm and illustrate the ZARU layout across the distal end of the q-arm on chromosome Z.

### Identification of chromosomal inversion in Silkie black-bone chicken

Following the annotation of ZARU at the distal end of chromosome Z, we examined structural variation between BBC and non-BBC breeds. Specifically, we examined the gapless CAU_Silkie and Huxu genome assemblies, focusing on potential structural rearrangements in this region. Pairwise alignment of these assemblies suggested the presence of an inversion at the distal end of chromosome Z.

We identified the genomic coordinates of each of the four ZARU genes on chromosome Z in both assemblies using blastn hits (see **Tables S2-S3**). Along with ZARU repeats, we also identified the tandem repeats across chromosome Z for both assemblies using the Global Repeat Map (GRM) application (Glunčić et al. 2022) (see **Table S4-S5**). Based on the identified ZARU gene repeats and tandem repeats, we constructed a map of repeat order for the distal end of chromosome Z.

Due to the presence of multiple copies of highly similar ZARU gene repeats, performing pairwise sequence alignment between the Huxu and Silkie assemblies in this region is challenging. To identify inversion boundaries, we compared the order of the repeat units, their lengths and orientations, and the relative distance between units of both assemblies. Through this manual comparison, we pinpointed rearranged junctions in Silkie BBC compared to the Huxu genome. We further validated this inversion in Silkie BBC by examining the junctions at each inversion boundary in detail. For Junction 1, we found a match spanning the initial segment of ZARU1 and the initial segment of inverted ZARU2; similarly, at Junction 2, we found a match spanning the inverted segment of ZARU1 with ZARU2. A comparison of repeat orders between the Huxu and Silkie assemblies enabled us to identify inversion boundaries at a higher resolution.

To confirm that the identified matches were not due to random occurrence, we used a custom Python script (available in our GitHub repository). The script examines ZARU unit order, length, and distances between consecutive units in both strands to identify matches, allowing for both exact and tolerance-based matches to allow variation in unit lengths and distance between units to accommodate divergence between the breeds. Finally, the script calculates a p-value using a binomial test based on exact matches. The matched occurrences are then organized and output with details such as direction, number of exact and tolerance-based matches, p-value, and interval values, presented in a sorted format for clearer interpretation.

### Phylogenetic, Principal component, and Admixture analysis

After validation of ZARU gene repeats distribution and inversion in Silkie BBC at the distal end of the q-arm of chromosome Z, we considered two single-copy regions, R1 (Z:78725000-80340000) and R2 (Z:81040000-82226500) which contains several protein-coding genes as potential *Id* locus regions. The R1 region is located entirely within the inverted region, while R2 has the same arrangement in BBC and non-BBC. The variant calls of the R1 and R2 regions was converted to phylip format using a custom Python script, vcf2phylip.py, downloaded from a GitHub repository (https://github.com/edgardomortiz/vcf2phylip). The resulting phylip file was then used as input for IQ-TREE 2 v2.3.4 (Minh et al. 2020) with -m MFP, --alrt 1000 -B 1000 -- boot-trees flags to construct localized phylogenies for both regions. To visualize the generated phylogenetic trees, we employed Figtree v1.4.2 (Rambaut 2014). Notably, we observed the separation of BBC from non-BBC in R1, while no such separation was seen in the R2 region.

In the case of the *Fm* locus, the phylogenetic tree of Dup1 and Dup2 regions separated the BBC from non-BBC individuals. The structural rearrangement at the *Fm* locus reduces the gene flow between BBC and non-BBC breeds, which gives rise to this phylogenetic separation. Therefore, we reasoned that the *Id* locus may also have the BBC-specific phylogenetic signal due to a chromosomal rearrangement. Since the *Id* locus boundaries are unknown, we constructed 10Kb non-overlapping sliding window-based phylogenies for the entire chromosome Z using IQ-TREE 2 with the same settings. In total, we obtained 8252 window-based trees. We filtered out 1003 trees marked as "failed" by IQ-TREE’s composition chi-square test due to significant deviations from average character composition, retaining only 7249 windows that passed the test. Visual interpretation from 7249 tree topologies is challenging. Therefore, we looked at the phylogenetic distance between these trees and the R1 topology. Using the 7249 qualified trees, we calculated the phylogenetic metrics distances between each of the 10Kb window trees and R1 region phylogeny using TreeCmp (Bogdanowicz et al. 2012). Specifically, five metrics (MatchingSplit, Robinson–Foulds, PathDifference, Quartet, and UMAST) suitable for unrooted phylogenies were utilized for the phylogenetic distance comparison. Finally, The results were visualized in R to explore the regions most closely associated with the R1 topology and are likely to be involved in the *Id* locus.

For the Principal Component Analysis (PCA) of both regions, we used PCAngsd v1.10 (Meisner and Albrechtsen 2018) based on genotype likelihood estimates derived from the Analysis of Next-Generation Sequencing Data (ANGSD) v0.935 (Korneliussen et al. 2014). The genotype likelihood values in beagle format, obtained from ANGSD, were employed in PCAngsd to determine principal components. Additionally, we estimated admixture proportions for both regions through NGSadmix (Skotte et al. 2013). In the NGSadmix analysis, we executed multiple runs with different values of K, ranging from K1 to K15, each with ten iterations. To discern the optimal K value, we applied the Evanno best K method (Evanno et al. 2005), an approach implemented in the Clumpak (Kopelman et al. 2015) web server, to identify the most appropriate K for our analysis. To visualize the results of PCA and Admixture analysis, we used R (Ihaka and Gentleman 1996).

### SNP Density Analysis

To estimate the SNP density across the entire chromosome Z, We employed a 10Kb non-overlapping sliding window-based approach. The common SNPs identified by both variant callers were used as input in VCFtools with the --SNPdensity 10000 flag. Due to the poor quality of variant calls in the ZARU regions, which contain repetitive sequences, we identified callable sites using the CallableLoci flag available in GATK v3.8.1 (McKenna et al. 2010). The results of the SNP density analysis of all sites and callable sites were then visualized using R.

### Genetic diversity and selection analysis

Our population genetic analysis exclusively included male (ZZ) individuals, a choice driven by the substantial genomic and sequence divergence observed in sex chromosomes. Using female (ZW) individuals can lead to uncertainty in the variant calls on the Z chromosome due to the mapping of gametologous or paralogous sequences shared between W and Z chromosomes (Ellegren 2011; Soh et al. 2014). Specifically, we selected three distinct breeds: two BBC breeds, Kadaknath (KADK) and Silkie (SILK), and a non-BBC breed represented by white leghorn chickens (WHLH). To ensure robust data quality, we carefully selected ten male individuals from each breed, each possessing sequencing coverage exceeding 20x (see **Table S6**). In pursuit of signatures of selection, we computed various population genetic statistics for the entire chromosome Z, employing 1 Kb windows as our analytical units. To assess genetic diversity within breeds, we utilized the folded site frequency spectrum (SFS) approach, implemented through ANGSD, to derive population-specific estimates of π (nucleotide diversity), Watterson’s θ, and τ (Tajima’s D). We calculated inter-population genomic differentiation (*F_ST_*) using ANGSD to quantify genetic differentiation between populations. We calculated the weighted *F_ST_* across chromosome Z by dividing the sum of the *F_ST_* value of the BBC vs non-BBC comparisons by the BBC vs BBC *F_S_*_T_ value.

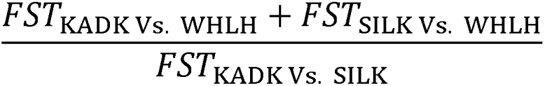

Additionally, we calculated divergence (*D_xy_*) between population pairs using the all-sites variant files, employing the popgenWindows Python script, which can be found at https://github.com/simonhmartin/genomics_general. To identify regions potentially under selection, we defined *F_ST_*, weighted *F_ST_*, and *D_xy_* outlier regions as 1 Kb windows falling within the top 10 percent of their respective estimates, signaling genomic regions of particular interest in population differentiation and evolutionary divergence.

### Identification of micro-RNA, their target genes, and gene enrichment analysis

Based on our criteria of identifying genomic regions under selection using population genetic analysis, we found a 26 Kb long (Z:72580000-72606000) intergenic region in the Pre-ZARU region. We searched for micro-RNAs within this region using a nucleotide sequence-based search of micro-RNAs using the miRBase (Kozomara and Griffiths-Jones 2014) webserver. Our search yielded the identification of five pre-micro-RNA precursors situated within the designated region. Subsequently, we sought to determine the target genes of these micro-RNAs. To accomplish this, we used the TargetScanHuman (Agarwal et al. 2015; McGeary et al. 2019) web server, specifying the chicken as the species of interest, and this enabled us to identify the target genes specific to these chicken-derived micro-RNAs. We performed gene enrichment analysis with the list of identified target genes. To explore the functional implications of these genes, we employed the KEGG pathway database (Kanehisa and Goto 2000) with default settings. This analysis was facilitated through the ShinyGO v 0.77 (Ge et al. 2020) web server, enabling us to gain insights into the potential pathways and biological processes influenced by these micro-RNAs and their associated target genes.

### Identification of BBC-specific SNPs

To understand the impact of genetic variants within both coding and non-coding regions, we used the snpEff v4.3t along with GRCg6a.96 databases for annotation (Cingolani et al. 2012). This approach allowed us to comprehensively assess the genetic variants within regions putatively under selection in BBC and promoter regions of various genes. Our analysis was precisely honed to focus on alternative SNPs found exclusively in BBC individuals while their non-BBC counterparts retained reference alleles. We employed the bam-read count method to calculate the proportions of SNPs within all 270 individuals (42 BBC and 228 non-BBC) (Khanna et al. 2022). Additionally, based on a nucleotide sequence-based search in PROMO (Farré et al. 2003) (a virtual laboratory for the identification of putative transcription factor binding sites) using a 5 % maximum matrix dissimilarity rate, we identified the transcription factors which bind on the promoter region of *CCDC112* which harbours 3 SNPs highly differentiated in BBC breeds.

## Results

### Resolution of ZARU and its units is better in GRCg6a than in GRCg7b assembly

In the telomere-to-telomere Huxu genome assembly, the ZARU regions are fully assembled without gaps. This completeness of the Huxu genome is further supported by our independent ONT long-read alignment, as displayed in the UCSC browser screenshot (**Fig. S1**), and by the Klumpy scan alignment (**Fig. S2**). Using the Huxu assembly as a reference for the distal end of the q-arm of chromosome Z, its comparison with GRCg6a and GRCg7b genome assemblies indicates that GRCg6a demonstrates higher assembly quality than GRCg7b (see **Fig. S3-S4**) in the distal end of chromosome Z. Specifically, ZARU2 is well-assembled in both the Huxu and GRCg6a assemblies, while in GRCg7b, ZARU2 appears to collapse with ZARU1 (see **Fig. S5-S6**). The Klumpy scan alignment further reveals that GRCg7b is more fragmented, containing more gaps relative to GRCg6a (see **Fig. S7-S8**). Although GRCg7b benefits from multi-platform, high-coverage data and exhibits generally high overall quality, its assembly of the ZARU regions remains suboptimal.

### *Id* locus is located between Z amplicon repeat units

Based on the read coverage calculation of the entire chromosome Z, we observed high coverage in some regions located at the end of chromosome Z (**Fig. S9**). Using the BLASTN and read coverage approaches, we have refined the precise positions of Z-amplicon repeat units (located at Z:73613150-82529921) situated at the distal end of the q-arm of chromosome Z within the GRCg6a genome (**Fig. 1**) and GRCg7b assembly (**Fig. S10**). We have identified two single-copy regions, R1 and R2, which contain several protein-coding genes and divide the Z amplicon repeat units into three distinct segments (ZARU1, ZARU2 and ZARU3). R1, spanning from Z:78725000 to Z:80340000, extends over 1.61 Mb and harbors ten protein-coding genes (*MTAP*, *CDKN2A*, *CDKN2B*, *TRIM36*, *PGGT1B*, *CCDC112*, *FEMC1*, *ALDH7A1*, *GRAMD2B*, and *ZNF608)*. Similarly, R2, spanning from Z:81040000 to Z:82226500, encompasses a length of 1.18 Mb and hosts more than 15 protein-coding genes. Previous research (Li et al. 2014; Tian et al. 2014b; Xu et al. 2017; Cha et al. 2023), particularly concerning dermal pigmentation and plumage color variation, has highlighted several candidate genes predominantly located within the R1 region. Considering these earlier studies and our findings (discussed in the subsequent section), it becomes evident that the *Id* locus is co-localized with the Z-amplicon repeat units at the distal end of the q-arm of chromosome Z.

**Fig. 1.**
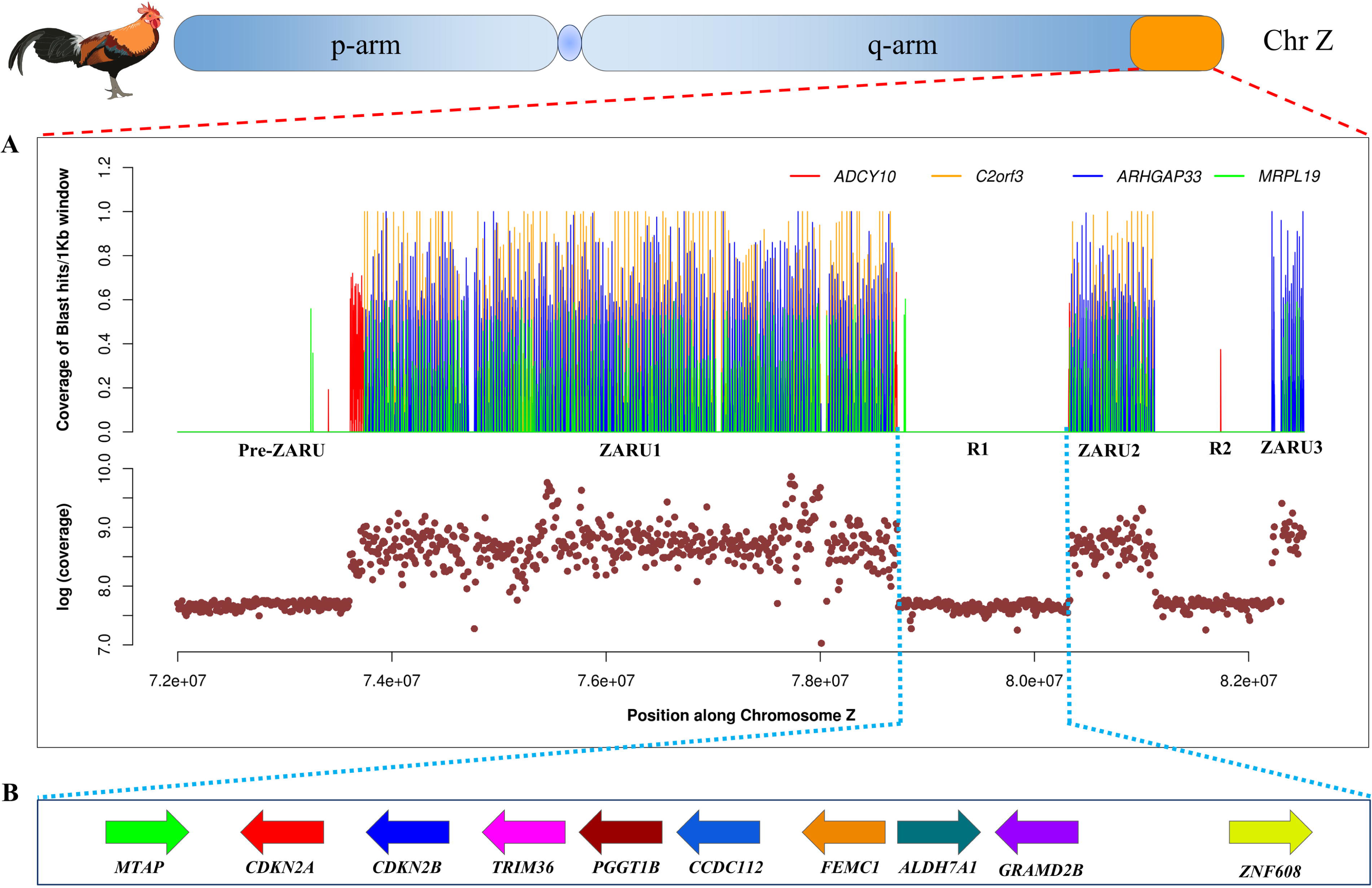
Magnified view of a ∼10.5 Mb segment (72,000,000–82,529,921 in GRCg6a genome assembly) on the distal end of the q-arm of chicken chromosome Z: **(A)** Detailed view of three sets of Z amplicon repeat units (ZARU) within this segment, containing multiple copies of the genes *C2orf3*, *ARHGAP33*, and *MRPL19 in* ZARU1 and ZARU2. Multiple copies of The *ADCY10* gene are present at the start and the end of ZARU1, while various copies of *ARHGAP33* and *MRPL19* are present in ZARU3, as illustrated in the top panel. The second panel displays the short-read coverage across this region in 10 Kb non-overlapping windows, highlighting elevated coverage due to gene repeats. **(B)** Schematic of the ten genes located in the R1 region, with arrows indicating the transcriptional direction of each gene.

### *Id* locus co-localizes with an inversion within ZARU in Silkie BBC

Through comparison of the organization of repeat units and tandem repeats on chromosome Z between the Huxu and Silkie assemblies, we observed that a substantial portion of ZARU1 (following breakpoint 1 at CP100594.1:75131357), along with the entire R1 region and the initial repeat units of ZARU2 (up to breakpoint 2 at CP100594.1:85496156), is inverted in Silkie BBC (see **Fig. 2A**). In Silkie, the initial segment of the inverted ZARU2 joins with the initial part of ZARU1 at junction 1 (see **Fig. 2B**). Similarly, the segment of ZARU1 following breakpoint 1 connects with ZARU2 at junction 2 (**Fig. 2B**).

**Fig. 2.**
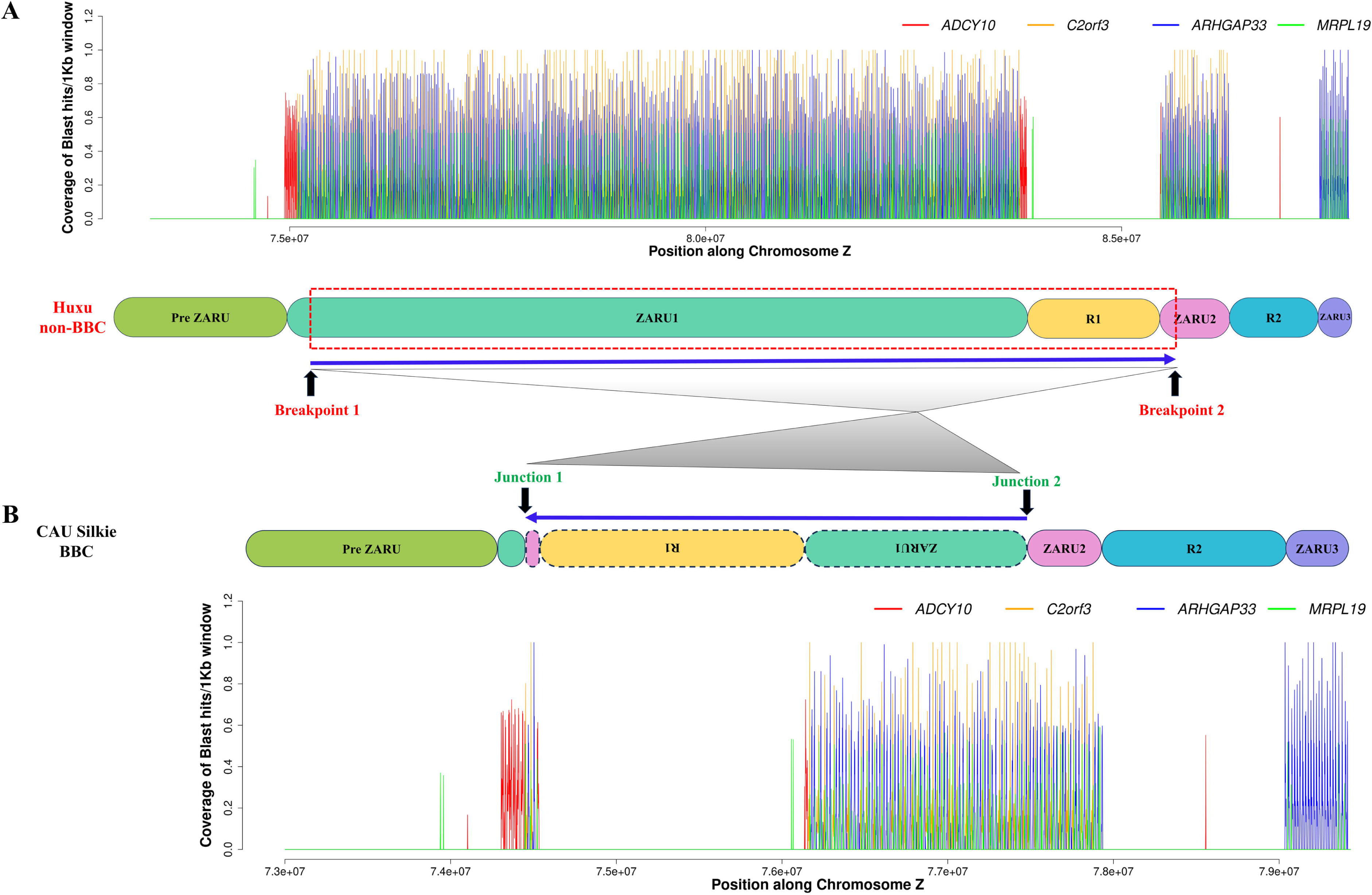
Chromosomal rearrangement at the distal end of the q-arm of chromosome Z in Silkie (BBC) compared to Huxu (non-BBC): **(A)** Schematic representation of three Z amplicon repeat units (ZARU1, ZARU2, and ZARU3), along with the Pre-ZARU, R1, and R2 regions at the distal end of chromosome Z. ZARU1 and ZARU2 contain multiple copies of the C2orf3, ARHGAP33, and MRPL19 genes. In ZARU1, the ADCY10 gene has multiple copies at both the beginning and end of the unit, while additional copies of ARHGAP33 and MRPL19 are found within ZARU3. The red, dotted rectangular box and breakpoints 1 and 2 highlight the genomic region, and breakpoints 1 and 2 highlight the genomic region inverted in the Silkie BBC genome. **(B)** Diagram of the chromosomal rearrangement in Silkie BBC, where the majority of ZARU1 (8.806 Mb of 8.97 Mb), the entirety of the R1 region (∼1.6 Mb), and the initial portion of ZARU2 (0.03 Mb) are inverted. This inversion results in a new configuration, with the initial portion of ZARU1 (0.191 Mb of 8.97 Mb) joined with the inverted segment of ZARU2 at Junction 1. The inverted ZARU1 is then joined with ZARU2 at Junction 2.

Along with the inversion, large regions of ZARU1 and ZARU2 were lost, resulting in a shorter ZARU region in Silkie BBC. A closer look at junction 1 reveals that the repeat order within ∼74Kb in Silkie (a match of 59 units) occurs in two segments in Huxu (the initial segments of ZARU1 (a match of 41 units) and ZARU2 (a match of 34 units) located ∼10.3Mb apart (see **Fig. 3** and **Table S7**). Likewise, the repeat order at junction 2, present within ∼72Kb in Silkie (a match of 70 units), occurs in two parts in Huxu (ZARU1 (a match of 69 units) after breakpoint 1 and ZARU2 (a match of 26 units)), positioned ∼10.6Mb apart (see **Fig. 4** and **Table S8**). These findings support the inversion of ZARU1 and the R1 region in Silkie BBC.

**Fig. 3.**
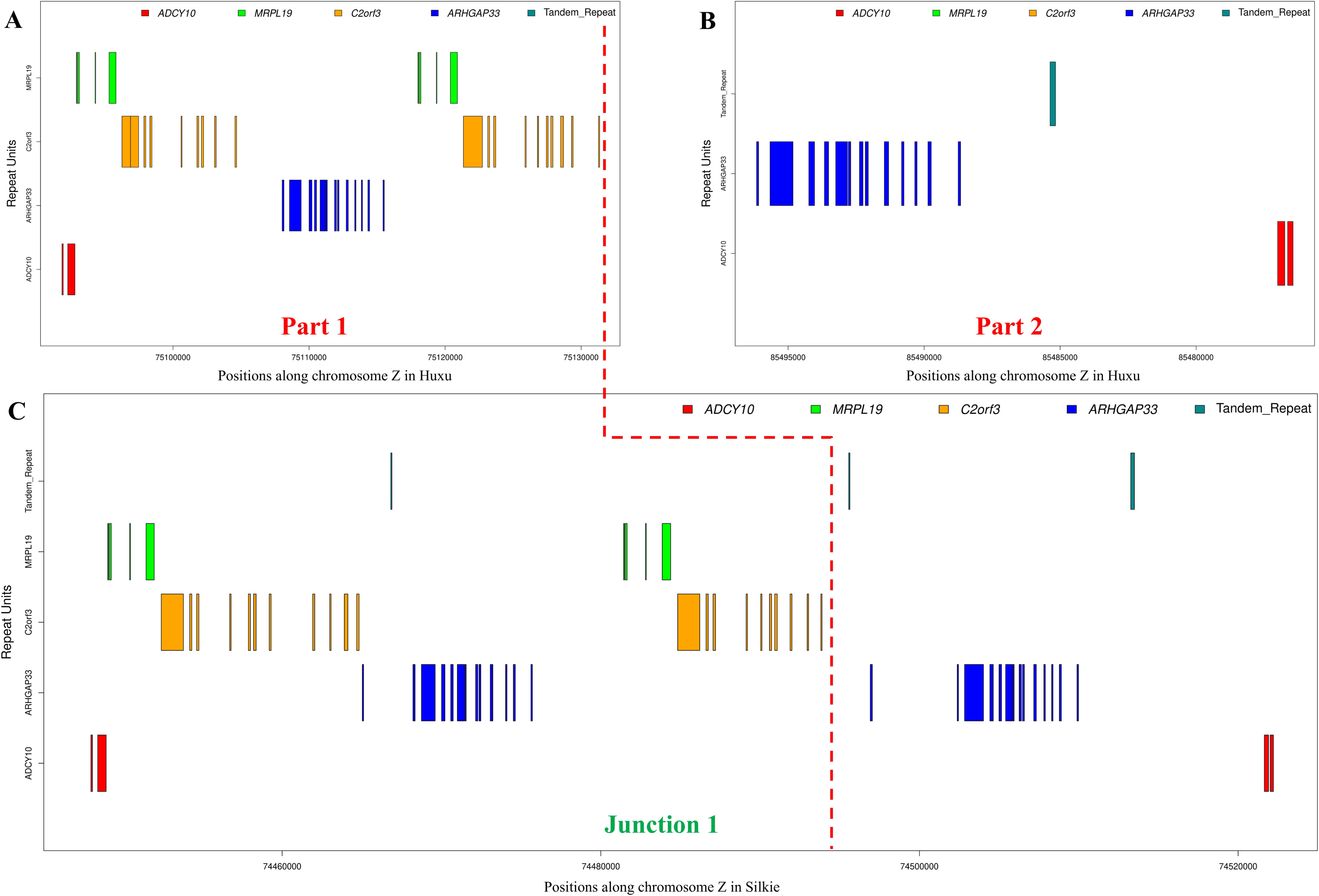
Zoomed View of Junction 1 in Silkie Black Bone Chicken (BBC): (A) and (B) illustrate the initial portions of ZARU1 and the inverted ZARU2, respectively, in the Huxu non- BBC genome. (C) shows Junction 1 in the Silkie BBC, where the initial segment of ZARU1 and the inverted segment of ZARU2 have fused. Exons are indicated by vertical rectangular boxes, color-coded by gene: *ADCY10* (red), *MRPL19* (green), *C2orf3* (orange), and *ARHGAP33* (blue). Tandem repeats are marked in dark cyan. The red vertical dotted line indicates the precise fusion point between the initial segment of ZARU1 and the inverted segment of ZARU2.

**Fig. 4:**
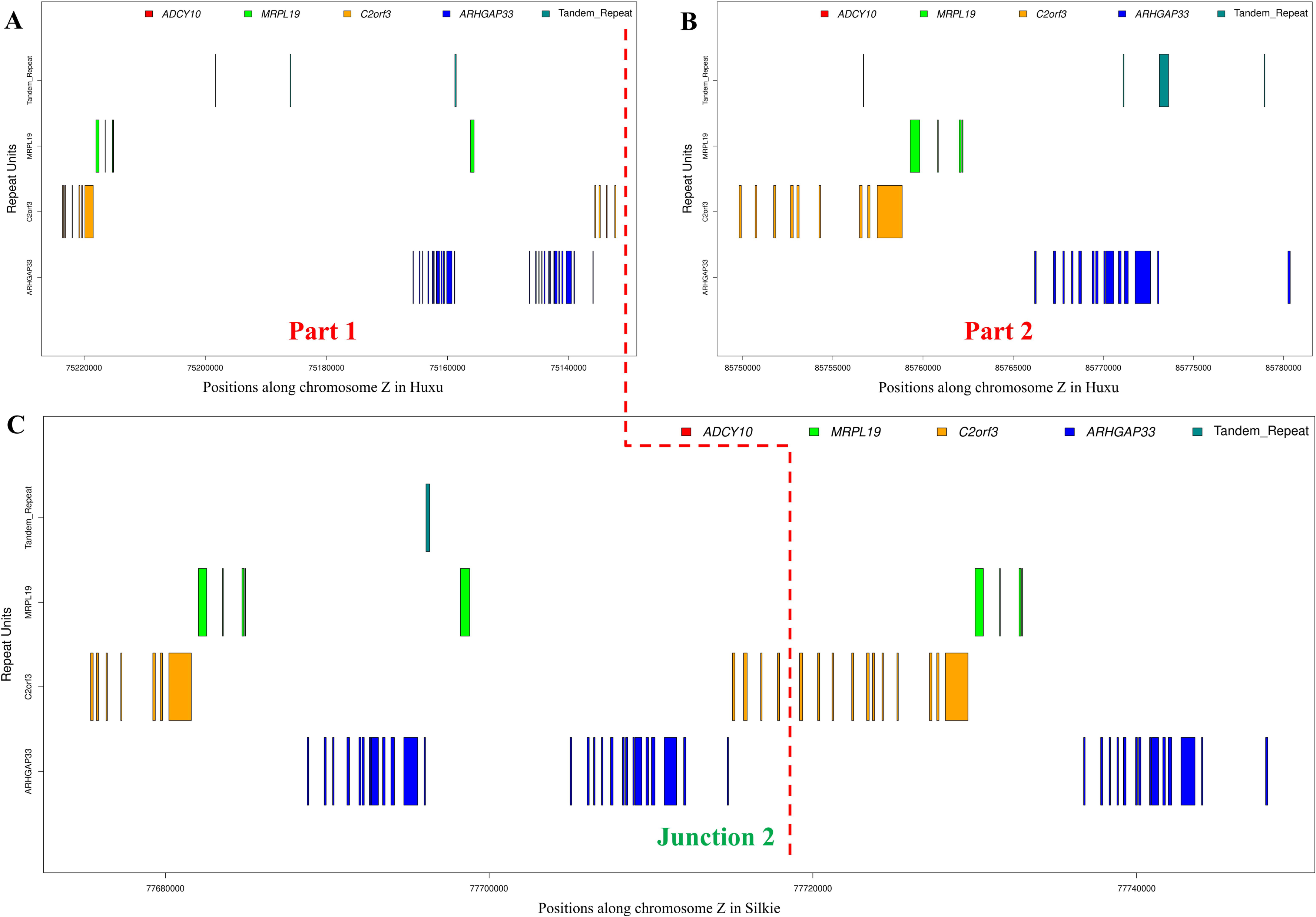
Zoomed View of Junction 2 in Silkie Black Bone Chicken (BBC): (A) and (B) illustrate the inverted ZARU1 and ZARU2, respectively, in the Huxu non-BBC genome. (C) shows Junction 2 in the Silkie BBC, where the inverted ZARU1 and the ZARU2 have fused. Exons are indicated by vertical rectangular boxes, color-coded by gene: *ADCY10* (red), *MRPL19* (green), *C2orf3* (orange), and *ARHGAP33* (blue). Tandem repeats are marked in dark cyan. The red vertical dotted line indicates the precise fusion point between the inverted ZARU1 and the ZARU2.

### BBC breeds share common ancestry at *Id* locus

Compared to the genome-wide phylogeny (see **Fig. S11**), the local phylogeny of the rearranged *Fm* locus separates the BBC breeds from non-BBC (see **Fig. S12**) due to the reduced gene flow. As we identified that the entire R1 and ZARU1 have been inverted in BBC breeds, *Id* locus may also have the BBC-specific separation due to this inversion. Our higher-resolution identification of the *Id* locus is based on a comprehensive analysis of local phylogeny, principal component analysis (PCA), and admixture analysis of the R1 and R2 regions. Our findings suggest that R1 represents the *Id* locus for the following reasons: The genetic structure of the R1 region demarcates BBC from non-BBC breeds (**Fig. S13-S20**). In stark contrast, when examining the R2 region, we observe an intermingling of BBC and non-BBC breeds (**Fig. S21-S25**). This intermixing implies a lack of distinct differentiation between these two groups in the R2 region. The local phylogeny analysis of the R1, based on single nucleotide polymorphisms (SNP) data from 270 individuals, provides clear insights into the genetic relationships among different chicken breeds. As illustrated in **Fig. 5A** and **Fig. S13**, this analysis largely separates BBC from non-BBC individuals. Notably, a few individuals from the Tibetan native chicken population exhibit genetic proximity to Tibetan BBC individuals, suggesting potential gene flow. The phylogenetic distance of the 10Kb window trees within R1 to the R1 topology is much less than the rest of the Z chromosome (see **Fig S14**). This pattern supports the conclusion that the entire R1 region possesses a distinct phylogenetic topology specific to BBC breeds.

**Fig. 5.**
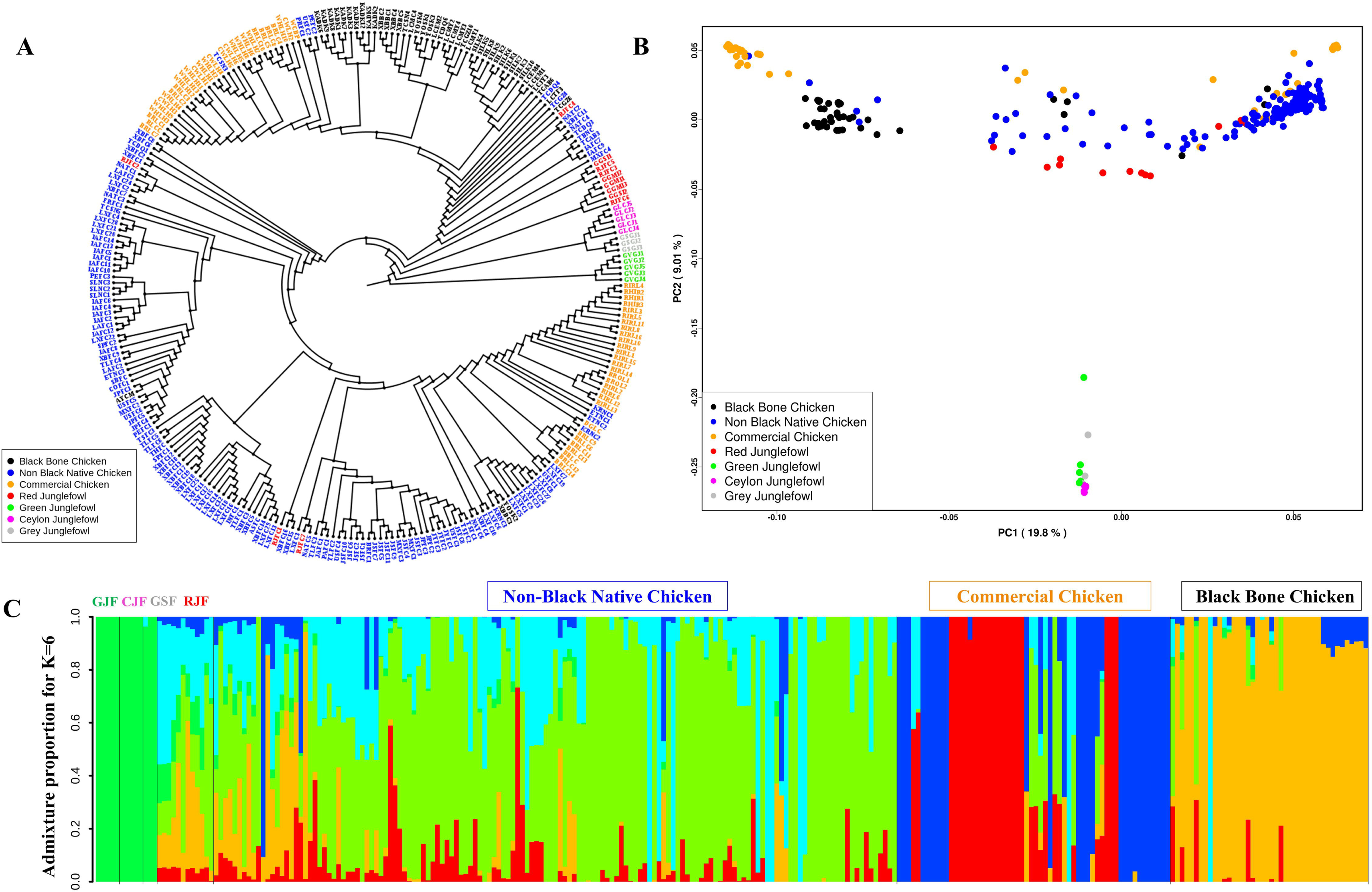
Phylogenetic and Population Structure Analysis of the R1 Region: **(A)** Phylogenetic tree based on the R1 region, including 270 individuals (42 Black Bone Chickens and 228 non-black chickens), generated in iqtree2 and visualized in FigTree to illustrate genetic relationships among sampled populations. **(B)** Principal Component Analysis (PCA) of the R1 region, where the first principal component (PC1) captures 19.8% of the total genetic variance and the second principal component (PC2) accounts for 9.01%, highlighting genetic differentiation across populations. **(C)** Admixture analysis at K=6 reveals distinct ancestry proportions, with all Black Bone Chickens forming a unique genetic cluster, indicating significant genetic separation from non-black chickens.

Principal component analysis (PCA) of the R1 region further elucidates the genetic structure, with PC1 and PC2 accounting for 19.8% and 9.1% of the genetic variation, respectively. Notably, most BBC individuals form a distinct cluster, reinforcing the patterns observed in the local phylogeny analysis (**Fig. 5B** and **Fig. S15-17)**. An admixture analysis of the R1 region, aimed at elucidating ancestry information, reveals that BBC individuals are most distinct when the best value of K (genetic clusters) is set to 6. Non-BBC native breeds exhibit limited genetic overlap with BBC and commercial breeds, as depicted in **Fig. 5C** and **Fig. S18-S20**.

### BBC-specific variants and genomic regions under selection

We employed two distinct approaches to find candidate genes or genomic regions associated with dermal pigmentation. In the first approach, we focused on identifying selective sweeps specific to BBC by leveraging various population genetic statistics. This analysis covered the distal end of the q-arm (Z:72000000-82529921). We sought to isolate the top 10% *F_ST_*windows, derived from pairwise comparisons between BBC and non-BBC breeds, as potential regions subject to selection. To designate a region as being under selection, we established specific criteria. *F_ST_* values were required to be less than 0.5 in comparisons involving BBC breeds, while in comparisons between BBC and non-BBC breeds, *F_ST_* values should exceed 0.8. Furthermore, the target region must fall within the top 10% of weighted *F_ST_* windows. This initial approach led us to identify numerous regions displaying substantial differentiation in BBC vs. non-BBC *F_ST_*comparisons while at the same time having lower differentiation in BBC vs. BBC *F_ST_* comparisons (**Fig. S26-28** and **Table S9** for detailed results).

In contrast to the signatures of selection, preexisting or standing genetic variation and breed-specific mutations arising from genetic drift can drive rapid evolution and influence phenotypic diversity and adaptation (Barrett and Schluter 2008). Consequently, our second approach investigates breed-specific genomic variants within BBC and non-BBC breeds. Our analysis has revealed the presence of numerous genomic variants in both genic and intergenic regions that are nearly fixed in both BBC and non-BBC breeds (**Table S10**).

### Putative BBC-specific selective sweep in the *MTAP* gene

Our genomic scan of R1 has unveiled distinct regions specific to BBC, characterized by a weighted *F_ST_* exceeding the predefined threshold. As depicted in **Fig. S26** and **Table S9**, one particularly noteworthy region spans five Kb and is situated within the last intron of the *MTAP* gene. What sets this 5 Kb region apart is its markedly lower *F_ST_*values in BBC vs. BBC comparisons compared to BBC vs. non-BBC comparisons, as evident in **Fig. 6A-D**. To further corroborate our findings, we examined *D_xy_* results within the same region, and they reinforced the observed *F_ST_* pattern, displaying striking differences, as depicted in **Fig. S29**. Within this 5 Kb region of the *MTAP* gene, we identified 6 BBC-specific SNPs, detailed in **Table S10**. Notably, the bam-read count data across 270 individuals underscore that these SNPs are nearly fixed among BBC individuals, irrespective of their geographic origins. A barplot depicting the percentage composition of each base at these six positions highlights the substantial difference in allele frequency between BBC and non-BBC individuals, as showcased in **Fig. 6E**.

**Fig. 6.**
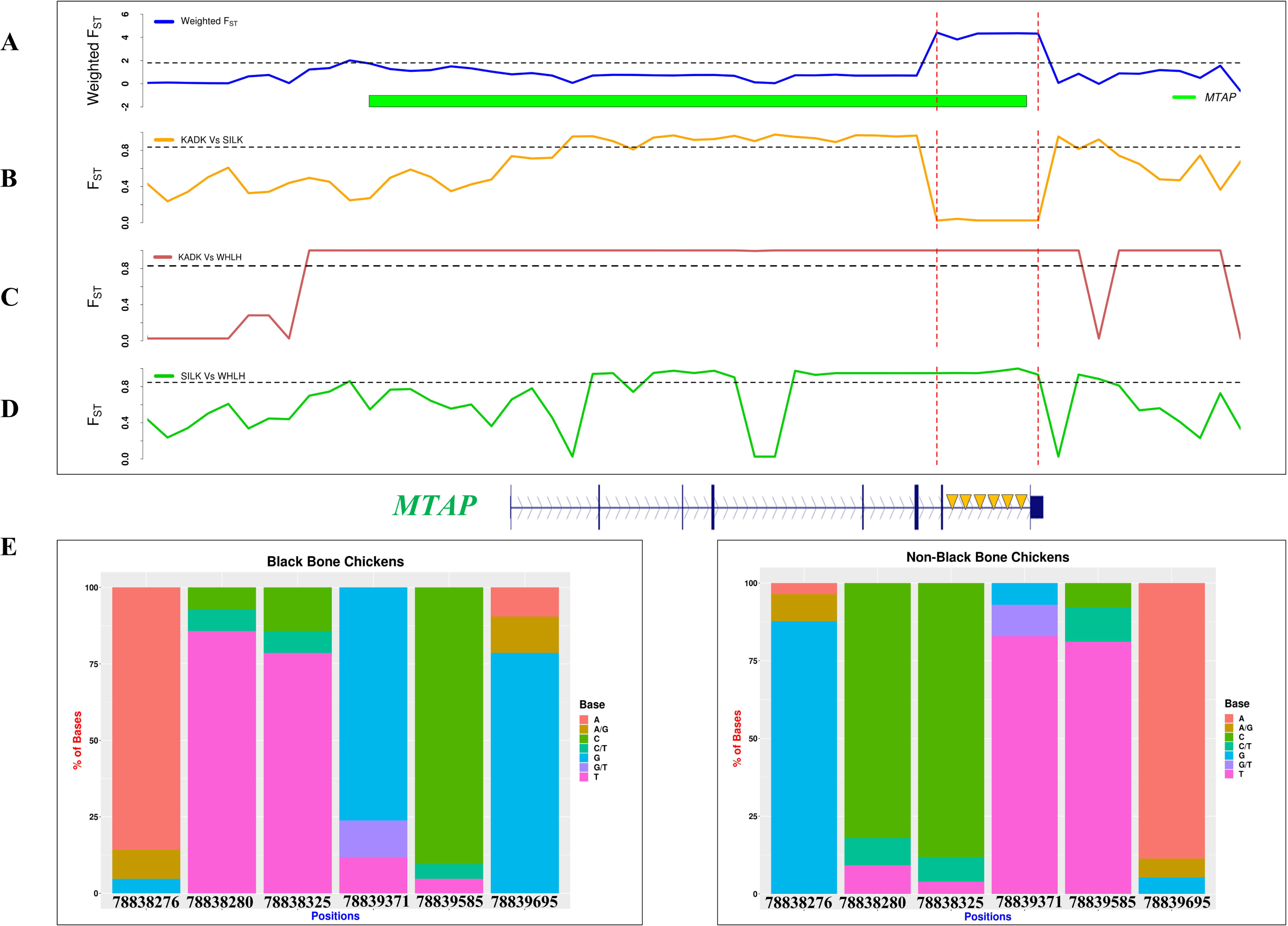
Signature of selection in the last intron of the *MTAP* gene within the R1 region: Comparative analysis of weighted *F_ST_* (calculated as the sum of BBC vs. non-BBC comparisons divided by BBC vs. BBC comparisons) and *F_ST_* values, using 1Kb non-overlapping windows. A horizontal black dotted line represents the 90th percentile threshold for both Weighted *F_ST_* and *F_ST_*values, highlighting regions of high differentiation. **(A)** Weighted *F_ST_*analysis comparing BBC (KADK and SILK) with non-BBC (WHLH) groups. **(B)** *F_ST_*comparison between KADK and SILK breeds. **(C)** *F_ST_* comparison between KADK and WHLH breeds. **(D)** *F_ST_* comparison between SILK and WHLH breeds. **(E)** Six BBC-specific SNPs with high differentiation within a 5 kb region in the last intron of the *MTAP* gene were identified. The percentage of bases at each SNP position is shown for both BBC (n=42) and non-BBC (n=228) breeds, indicating breed-specific allele frequencies.

### BBC-specific variants in the promoter region of the *CCDC112* gene

Within the promoter region of the *CCDC112* gene, we identified three highly differentiated SNPs located at positions Z:79049203, Z:79049207, and Z:79049235. Our analysis across 42 BBC individuals found that ∼83% are homozygous for the alternative allele, ∼9% are heterozygous, and ∼7% possess the reference allele. To provide a more accessible and insightful visualization of these findings, we’ve generated a barplot illustrating the bam-read count results for these three positions, as showcased in **Fig. 7**. In the case of the 228 non-BBC individuals, ∼72% have reference alleles, ∼12% are heterozygous, and ∼ 14% possess alternative alleles, akin to those observed in BBC individuals. These large differences in allele frequency underscore the genetic divergence in this gene between BBC and non-BBC breeds. These three highly differentiated BBC-specific SNPs located within the promoter region of *CCDC112* overlap with binding sites of 27 transcription factors (see **Table S11**).

**Figure 7.**
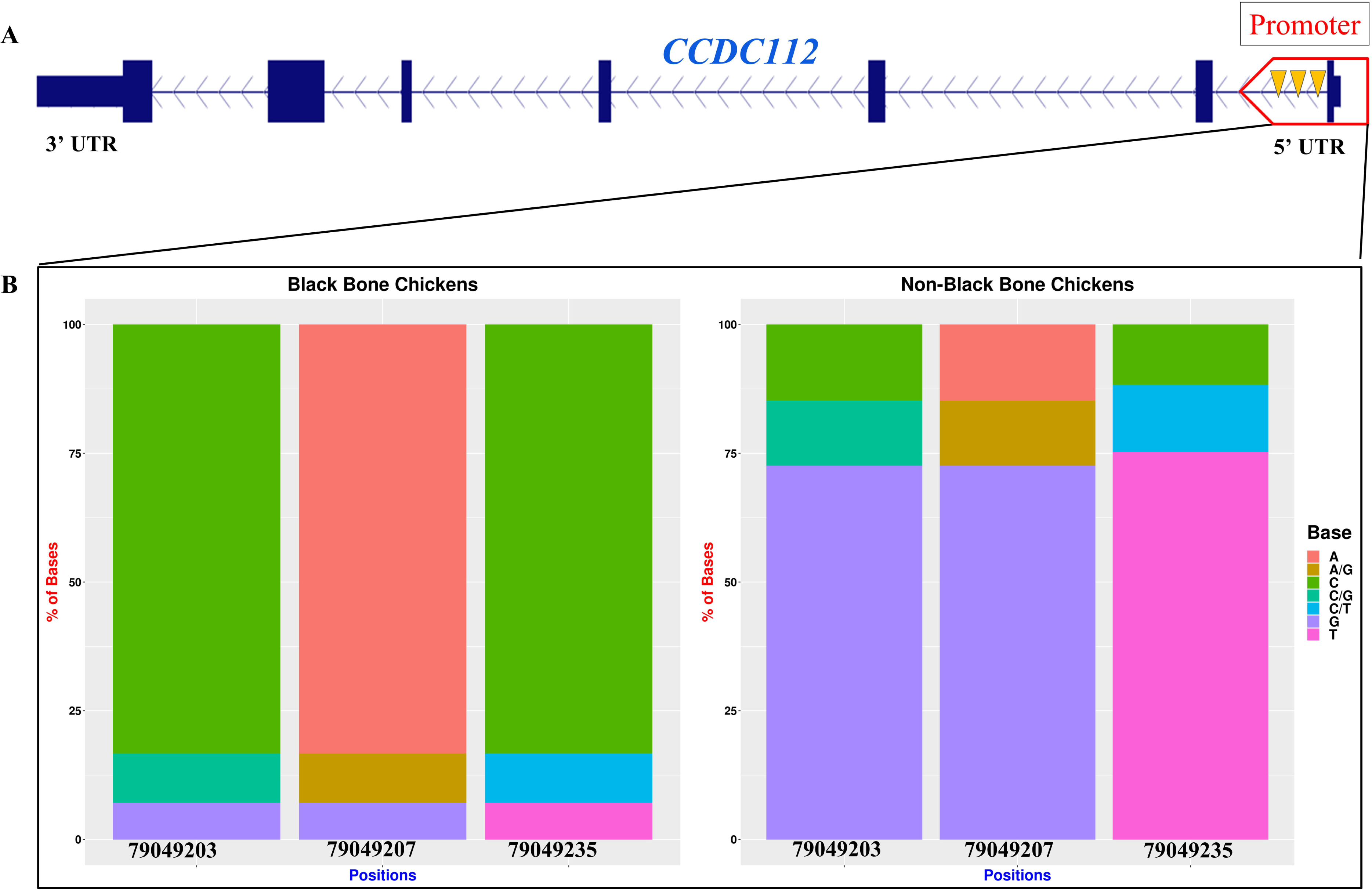
Black-bone chicken-specific SNPs in the promoter region of the *CCDC112* gene: **(A)** Illustration of three distinctive SNPs located within the promoter region of the *CCDC112* gene, unique to BBC, marked by orange triangles. **(B)** Allele frequency analysis shows the percentage of bases at each SNP position for the BBC breeds (n=42) and the non-BBC breeds (n=228), highlighting breed-specific genetic variation.

### High differentiation of BBC-specific variants in R1 compared to the rest of chromosome Z

The ZARU regions on chromosome Z show a unique SNP distribution, likely influenced by poor short-read mapping quality, which can increase false SNP calls (see **Fig. S30**). Although SNP density in callable sites within ZARU regions is slightly lower than the rest of chromosome Z, the difference is minimal. Specifically, the R1 region exhibits BBC breed-specific variants with an SNP density of 0.043/base, similar to the overall chromosome Z density of 0.040/base.

In R1, we identified nine highly differentiated SNPs, six of which were in the *MTAP* gene and three in the *CCDC112* gene. This frequency of differentiated SNPs in R1 (0.0132%) notably exceeds the expected chrZ-wide proportion (0.00288%), where only 1.97 differentiated SNPs would be anticipated if distributed randomly. A binomial test confirmed the statistical significance of this excess, yielding a p-value of 0.00021, indicating that the observed concentration of differentiated SNPs in R1 is unlikely to be due to chance.

### Pre-microRNA precursors identified in BBC-specific region target melanogenesis genes

We identified several other genomic regions in the Pre-ZARU region upstream with a selection signature in BBC breeds (**Table S9**). Among these, we found a 26 Kb continuous intergenic region spanning Z:72580000-72606000 with a remarkable pattern in pairwise *F_ST_* and *D_xy_* comparisons, as depicted in **Fig. 8A-D**. The examined *D_xy_* results within the same region supported the observed *F_ST_* pattern, displaying striking differences (see **Fig. S31**). The nucleotide diversity (π) within this region is higher in White Leghorn chickens. However, Kadaknath and Silkie chickens have markedly reduced nucleotide diversity, as visualized in **Fig. S32**.

**Fig. 8:**
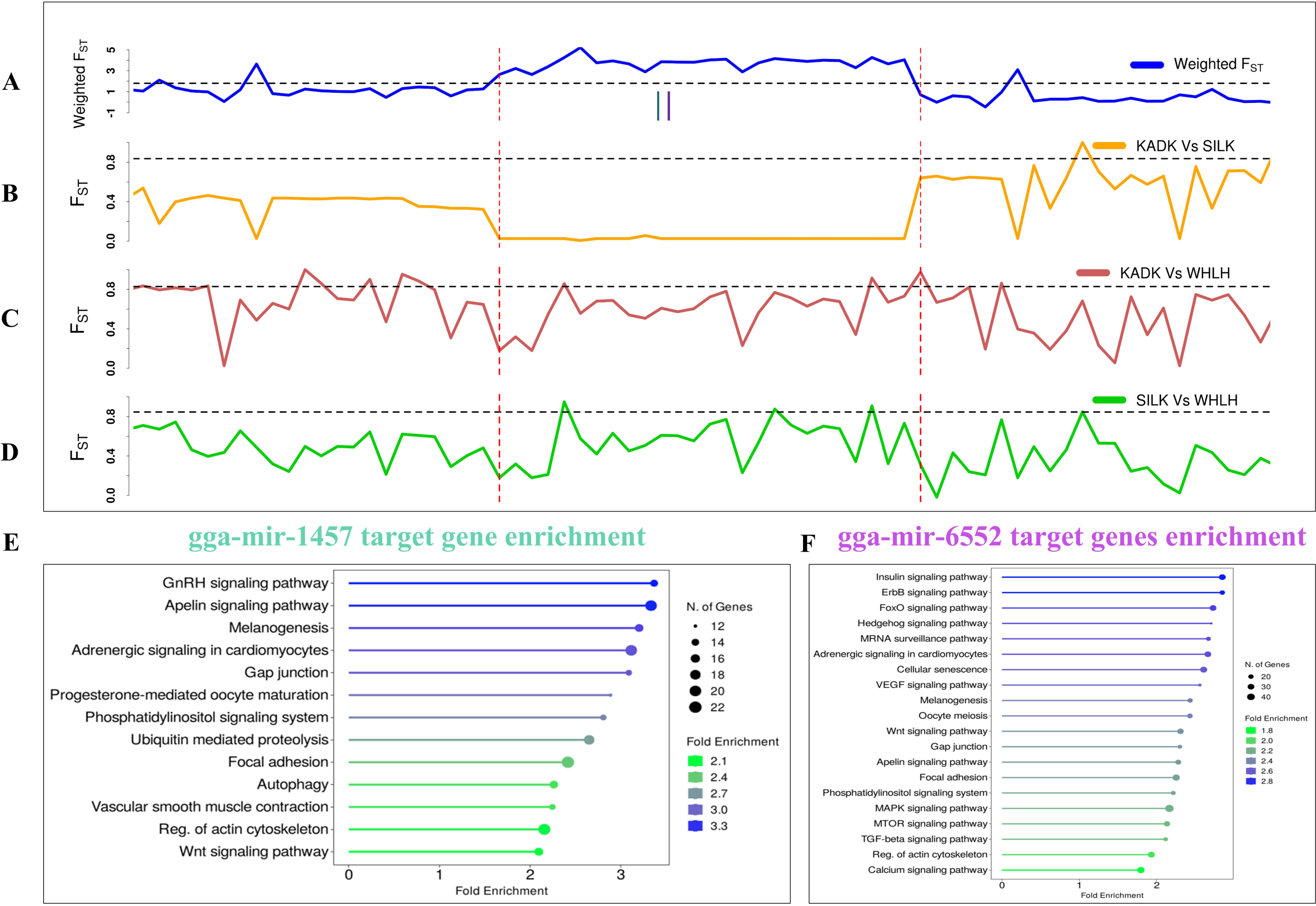
Signature of selection in a 26 Kb long non-geneic region located in the Pre-ZARU region: Comparative analysis of weighted *F_ST_* (calculated as the sum of BBC vs. non-BBC comparisons divided by BBC vs. BBC comparisons) and *F_ST_* values, using 1Kb non-overlapping windows. A horizontal black dotted line represents the 90th percentile threshold for both Weighted *F_ST_* and *F_ST_*values, highlighting regions of high differentiation. **(A)** Weighted *F_ST_*analysis comparing BBC (KADK and SILK) with non-BBC (WHLH) groups. **(B)** *F_ST_*comparison between KADK and SILK breeds. **(C)** *F_ST_* comparison between KADK and WHLH breeds. **(D)** *F_ST_* comparison between SILK and WHLH breeds. **(E)** and **(F)** presents the gene enrichment analysis for the target genes of gga-mir-1457 and gga-mir-6552.

Although this specific region lacks any protein-coding genes, we uncovered five pre-microRNA precursors highly similar to gga-mir-1457, gga-mir-6702, gga-mir-6551, gga-mir-6555, and gga-mir-6552 (**Fig. S33**). Of these five miRNAs, we found distinct BBC-specific SNPs within the paralogs of gga-mir-1457 and gga-mir-6552, which are predicted to target several genes (**Table S12-13**). Gene enrichment analysis of the targets of these two miRNAs revealed multiple target genes associated with various signaling pathways, including the GnRH signaling pathway, Apelin signaling pathway, Insulin signaling pathway, adrenergic signaling pathway, Wnt signaling pathway, and Hedgehog signaling pathway (**Fig. 8E-F**, and **Fig. S34-35**). Notably, our analysis in the context of gene enrichment unveiled an intriguing finding: 30 genes integral to the melanogenesis pathway (see **Table S14)** serve as targets for these miRNAs, underscoring their potential role in regulating pigmentation-related processes.

## Discussion

Our study has refined the position of the *Id* locus within a large (∼10.36 Mb) inversion and identified several candidate *Id* locus genes. We demonstrate that the *Id* locus is co-localized among Z amplicon repeats, and the causative mutations likely occur within a region of ∼1.61 Mb (i.e., R1) at the distal end of the q-arm of chromosome Z. The phylogenetic and population structure analysis of 270 individuals (42 BBC and 228 non-BBC) suggest that the *Id* locus has common ancestry among all BBC breeds. Within the refined region of the *Id* locus, we identified the BBC-specific regions putatively under selection. The two candidate genes, *MTAP* and *CCDC112*, involved in the melanogenesis pathway and plumage color variation, have BBC-specific SNPs in the intronic and promoter regions, respectively. The findings of our study suggest that the *Id* locus has a common ancestral state in BBC breeds and a polygenic nature, which is responsible for the variation in dermal pigmentation.

Chromosomal inversions are key in evolution, often suppressing recombination, limiting gene flow, and enabling the accumulation of breed or population-specific mutations, which contribute to developmental, morphological, and phenotypic diversity (Kirkpatrick and Barton 2006; Wellenreuther and Bernatchez 2018; Recuerda and Campagna 2024). Large inversions have been documented in birds, such as a 115 Mb inversion in quail (Sanchez-Donoso et al. 2022), four inversions ranging from 12-63 Mb in the zebra finch (Knief et al. 2016), and a 4.5 Mb-long inversion in the ruff (Küpper et al. 2015; Lamichhaney et al. 2015) linked with morphological variation, fitness and reproductive strategies. In chicken, a 7.4 Mb long inversion at chromosome 7 has been reported, which causes a rose-comb phenotype and poor sperm motility (Imsland et al. 2012; Wang et al. 2017). Overall, the role of chromosomal rearrangements in complex phenotypes is increasingly being recognised with the availability of genomic tools.

Our study has identified one of the longest novel chromosomal inversions (∼10.36 Mb) at the highly repetitive distal end of chromosome Z in Silkie BBC. Our novel approach of using ZARU genes and tandem repeats as markers has shown that the *Id* locus is co-localized within Z amplicon repeat units and has undergone an inversion. This inversion could harbor genetic changes causing inhibition of dermal pigmentation in BBC chickens. Identifying the causal mutation is challenging in such cases when a large inversion containing several genes is associated with a complex phenotype. Many sequence polymorphisms in strong linkage show a substantial association with the phenotype. Moreover, it is possible that multiple genes or multiple SNPs associated with the same gene could be responsible for the inhibition of dermal pigmentation. Since *Id* is sex-linked and has incomplete dominance, additional mutations contributing to the same inhibition phenotype could have accumulated within the inversion. Future studies relying on high-quality genome assemblies and comprehensive annotation of this region will be crucial for obtaining definitive answers.

The hyperpigmentation phenotype in the BBC breeds results from common complex chromosomal rearrangement at the *Fm* locus and distinguishes them from non-BBC breeds (Dharmayanthi et al. 2017; Shinde et al. 2023). The shared rearrangement in diverse BBC breeds supports a model of a common origin of BBC followed by dispersal globally. Therefore, we anticipated that the *Id* locus should possess unique characteristics specific to and shared among BBC breeds. Our focus on R1 as a potential *Id* locus led us to perform local phylogenetic analysis, principal component analysis (PCA), and admixture analysis. These analyses effectively separated the BBC breeds from the non-BBC breeds, emphasizing the common ancestry of the *Id* locus within the BBC breeds similar to the *Fm* locus. We observed relatedness at the *Id* locus between some BBC and Tibetan non-BBC native breeds, suggesting ongoing gene flow between these groups reported in earlier studies (Li et al. 2020). Hence, our analysis indicates that the *Id* locus emerged in BBC before their global dispersal and may share selection signatures and causative genetic variants involved in inhibiting dermal pigmentation.

One of our major findings is the signature of selection in the 5Kb genomic region of the last intron of *MTAP,* which contains 6 SNPs with an exceptionally high percentage of BBC-specific fixation. *MTAP* plays a crucial role in the melanogenesis pathway within melanocytes, the specialized cells responsible for producing the melanin pigment in the skin (Ainger et al. 2017). It is responsible for catalyzing the phosphorylation of methylthioadenosine (MTA), a byproduct of polyamine metabolism (Behrmann et al. 2003). MTA, in turn, hinders the function of phosphodiesterase (PDE), a regulator of intracellular cAMP levels. By blocking PDE, MTA blocks the cAMP-PKA-CREB pathway, leading to the inhibition of *MITF* stimulation. It has been reported that mutations in the *MTAP* gene are associated with nevus count and melanoma development (Kvaskoff et al. 2011). Genomic variants identified in the *MTAP* are associated with plumage and dermal pigmentation in the F2 population crossed from White Leghorn and the Yeonsan Ogye (Heo et al. 2023; Cha et al. 2023). Functional gene expression analysis of Silkie and White Leghorn chicken embryos during early embryonic stages indicated distinct patterns of *MTAP* expression. Before embryonic day 10, White Leghorn embryos exhibited a higher *MTAP* expression level than Silkie embryos. However, a shift occurred after day 10, with Silkie embryos showing greater *MTAP* expression than White Leghorn embryos (Tian et al. 2014b). In the context of migration and differentiation studies of melanoblasts in Silkie and White Leghorn chickens, it was observed that Silkie neural crest cells underwent a slower melanocyte differentiation process compared to their White Leghorn counterparts. Specifically, on day 8, Silkie neural crest cells displayed less pigmentation than their White Leghorn counterparts. However, by day 11, they had achieved full pigmentation, signifying a distinct temporal pattern in melanocyte development between these two chicken breeds (Reedy et al. 1998; Faraco et al. 2001). The expression of *MTAP* and the migration and differentiation of melanocytes, both displaying a similar pattern, strongly imply the pivotal role of *MTAP* in regulating dermal pigmentation.

The sex-linked loci barring (B), dermal melanin inhibitor (*Id*), Translocational breakpoint (TB) and recessive white skin (y) are reported to occur in a linear order as B-*Id*-TB-y (Bitgood 1985, 1988). Subsequently, the sex-linked barring locus (B) was mapped to the *CDKN2A*/*2B* gene in the Barred Plymouth Rock chicken breed (Hellström et al. 2010). The *MTAP* gene occurs earlier (towards the centromere) than *CDKN2A/2B* in non-BBC breeds. Hence, the genomic position of the *MTAP* gene does not match with the B-*Id*-TB-y linear ordering of loci. However, after the inversion of the entire R1 region along with ZARU1, *CDKN2A/2B* is now located earlier than *MTAP* in BBC breeds. The rearrangement establishes the linear relationship of B-*Id*-TB-y, where *CDKN2A/2B* represents B, and *MTAP* can represent *Id*. Our findings of the R1 spanning inversion and putative selective sweep with SNPs suggest that *MTAP* is the most prominent candidate gene for the dermal melanin inhibitor.

SNPs in gene promoters can have implications for traits such as skin color and plumage in various species. For instance, in humans, an SNP in the promoter of the *SLC45A2* gene has been linked to variations in skin color (Graf et al. 2007). In ducks, the black plumage phenotype is determined by a mutation within the regulatory region of the *MC1R* gene (Liu et al. 2023). Similarly, studies on different chicken breeds have revealed that mutations in the promoter regions of genes like *MITF*, *TYR*, *ASIP,* and *GJA5* play a role in influencing skin color (Yu et al. 2017, 2019; Wang et al. 2018; Li et al. 2021). Interestingly, past research has established a connection between loss-of-function mutations in *CCDC112* and variations in feather color in geese (Ren et al. 2021). A previous gene expression analysis study conducted between Silkie BBC and White Leghorn chickens reported that after embryonic day 10, the *CCDC112* expression within the shank of Silkie embryos was greater than that of White Leghorn embryos (Tian et al. 2014b). This finding strongly implies a potential role of *CCDC112* in shank pigmentation in chickens. The regulatory mutations identified within *CCDC112’s* promoter region hold promise for shedding light on their association with dermal pigmentation and plumage color. The transcription factors with binding sites overlapping three highly differentiated promoter SNPs in BBC breeds are primarily involved in developmental patterning (Lorenzo et al. 2017; Minto et al. 2024), cell differentiation (Darlington et al. 1998), immune response regulation (Tsukada et al. 2011; Villarino et al. 2015), and sex determination (Larney et al. 2014).

We found that paralogs of two miRNAs, gga-mir-1457 and gga-mir-6552, which have BBC-specific SNPs, target 30 genes associated with the melanogenesis pathway. Previous literature has extensively documented the role of miRNAs in melanocyte biology and pigmentation, primarily by modulating the expression of specific genes or pathways (Mione and Bosserhoff 2015). Associations between miRNAs and melanin production, skin pigmentation, and plumage coloration have been observed in diverse species such as Koi fishes (Tian et al. 2018), pigs (Yuan et al. 2023), alpacas (Zhu et al. 2019) and ducks (Apopo et al. 2015). Numerous miRNAs have been identified in the case of Muchuan BBC, impacting melanin production in the breast muscle (Yu et al. 2020). The miRNA in the pre-ZARU may be involved in plumage color regulation and require a comprehensive functional analysis, which can provide valuable insights into their roles within the melanogenesis pathway.

The Z Amplicon Repeat Units (ZARU) situated at the distal end of chromosome Z harbor hundreds of copies of protein-coding genes, including *ADCY10*, *C2orf3*, *ARHGAP33*, and *MRPL19*. Despite their abundance, the exact function of ZARU remains elusive, leaving open the possibility of additional structural rearrangements within this region that may be involved in complex epigenetic gene regulation and its potential association with the inhibition of dermal melanin production. For instance, the tandem repeats within the ZARU region are known to have an intricate methylation pattern (Eden et al. 1981). The repetitive nature of this region also poses significant challenges for mapping short reads and variant calling, which often leads to erroneous results (DePristo et al. 2011; Koboldt 2020). Consequently, analyzing the ZARU region under current technical conditions is challenging. Nonetheless, the availability of long-read data, advancements in mapping and variant calling pipelines hold promise for unraveling the mysteries surrounding this region in the future.

## Conclusion

We explored the *Id* locus using read coverage analysis and blastn search for annotating amplicon repeats to map the organization of ZARU and localized population genetic structure analysis to refine its position to identify candidate genes. Our investigation concludes that the *Id* locus (Z:78725000-80340000) closely co-localizes with ZARU situated at the distal end of the q-arm of chromosome Z with a BBC-specific inversion. Multiple lines of evidence, such as local phylogeny, PCA, admixture and highly differentiated genomic variants, support the common ancestry of the R1 region within BBC breeds similar to the *Fm* locus. The population genetic signatures, the genome assembly of CAU_Silkie and the relative organisation of ZARU spanning the R1 region are consistent with a large chromosomal inversion in all BBC breeds. Among the identified genes, *MTAP* is the most prominent candidate for the *Id*, suggesting a pivotal role in inhibiting dermal pigmentation. In addition, *CCDC112* emerges as a potential candidate for influencing variations in pigmentation patterns. Further scrutiny of BBC breeds-specific selective sweep region in the Pre-ZARU region led us to discover miRNAs that can target various genes involved in the melanogenesis pathway. Our findings collectively suggest that the *Id* locus is potentially polygenic and strongly associated with the inversion with shared ancestry among BBC breeds.

## Availability of data

Scripts and data are available at https://github.com/Ashu2195/ID_locus.

## CRediT authorship contribution statement

**Ashutosh Sharma:** Conceptualization, Formal analysis, Investigation, Visualization, Validation, Writing - original draft, Writing - review & editing. **Nagarjun Vijay:** Conceptualization, Resources, Writing - original draft, Writing - review & editing, Funding acquisition, Project administration, Supervision.

## Acknowledgment

We thank the University Grants Commission for supporting AS with a Ph.D. scholarship. The Department of Biotechnology, Ministry of Science and Technology, India (Grant no. BT/11/IYBA/2018/03) and Science and Engineering Research Board (Grant no. ECR/2017/001430) provided funds used to generate primary sequencing data published in this article and computational resources (i.e., Har Gobind Khorana Computational Biology cluster) used.

## Appendix

Supplementary 1. Supplementary figures Supplementary 2. Supplementary tables

## Notes

https://github.com/Ashu2195/ID_locus

### Competing Interest Statement

The authors have declared no competing interest.

### Summary of Updates

The manuscript has additional analysis to refine the location of the Id locus. Importantly, using new genome assemblies, we show that Id locus overlaps a large chromosomal inversion in the ZARU region.

