## Supplementary Figures for "Common Ancestry of the *Id* Locus: Chromosomal Rearrangement and Polygenic Possibilities"

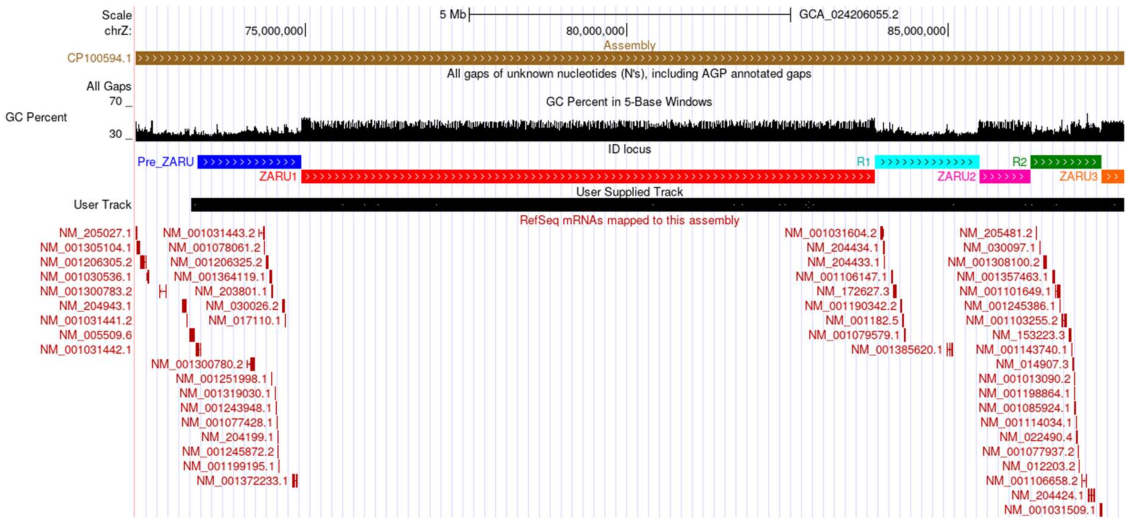


**Fig. S1: Evaluation of Huxu genome assembly using ONT long reads:** Screenshot of the UCSC genome browser showing the alignment of long reads (SRR15421342, SRR15421343, SRR15421344, SRR15421345, SRR15421346, SRR13494713, and SRR13494714) generated using ONT technology and aligned to the Huxu genome using minimap2.

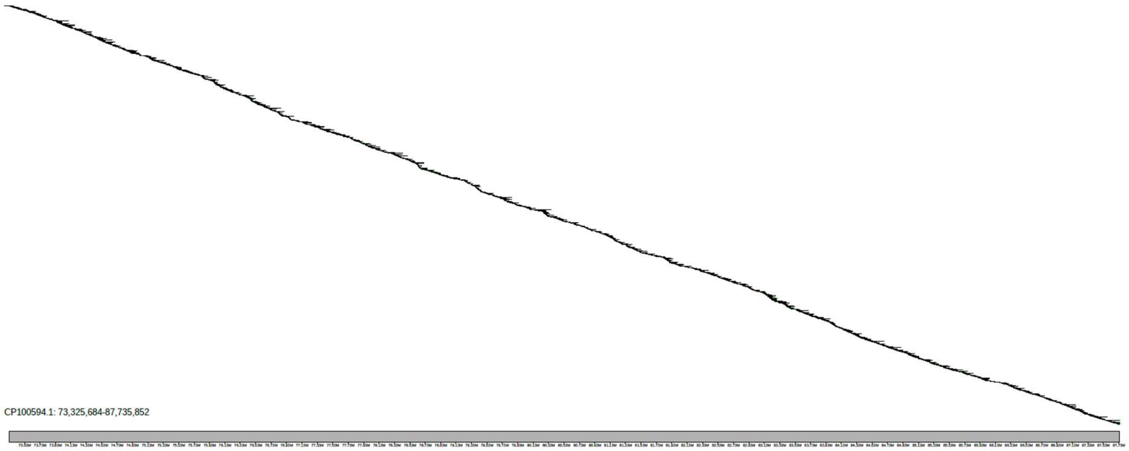


**Fig. S2: Evaluation of Huxu genome assembly using ONT long reads:** Klumpy scan alignment plot of ONT long-reads (SRR15421342, SRR15421343, SRR15421344, SRR15421345, SRR15421346, SRR13494713, and SRR13494714) aligned to the Huxu genome assembly using minimap2.

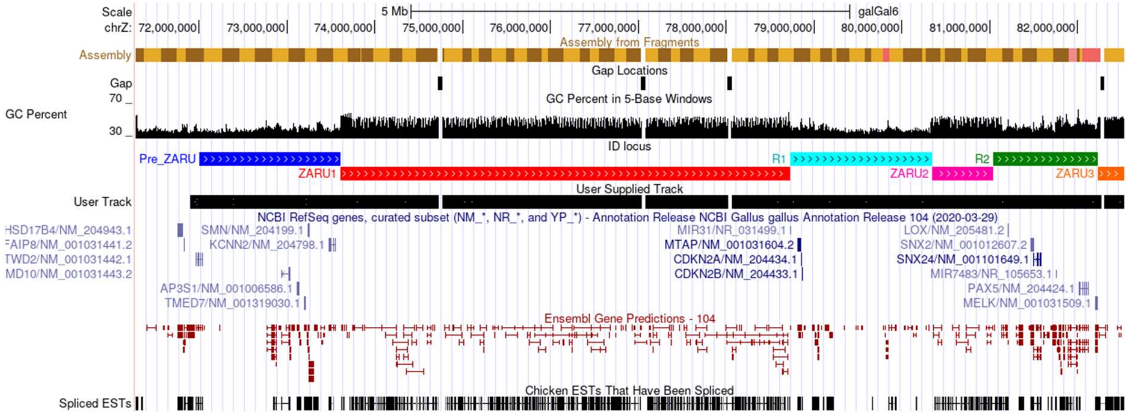


**Fig. S3: Evaluation of GRCg6a genome assembly using ONT long reads:** Screenshot of the UCSC genome browser showing the alignment of long reads (SRR15421342, SRR15421343, SRR15421344, SRR15421345, SRR15421346, SRR13494713, and SRR13494714) generated using ONT technology and aligned to the GRCg6a genome using minimap2.

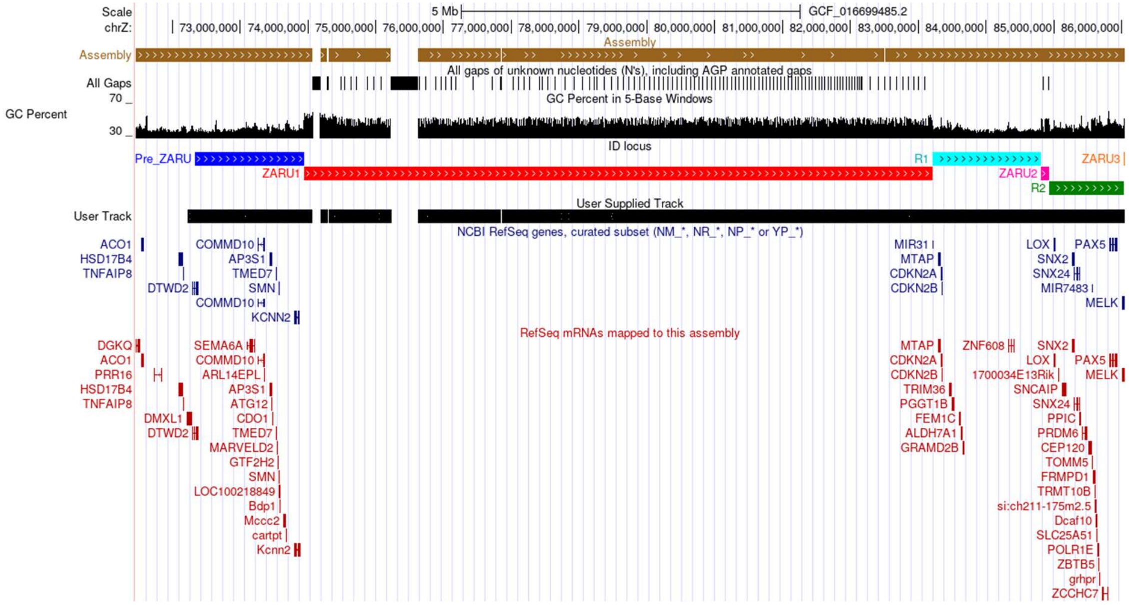


**Fig. S4: Evaluation of GRCg7b genome assembly using ONT long reads:** Screenshot of the UCSC genome browser showing the alignment of long reads (SRR15421342, SRR15421343, SRR15421344, SRR15421345, SRR15421346, SRR13494713, and SRR13494714) generated using ONT technology and aligned to the GRCg7b genome using minimap2.

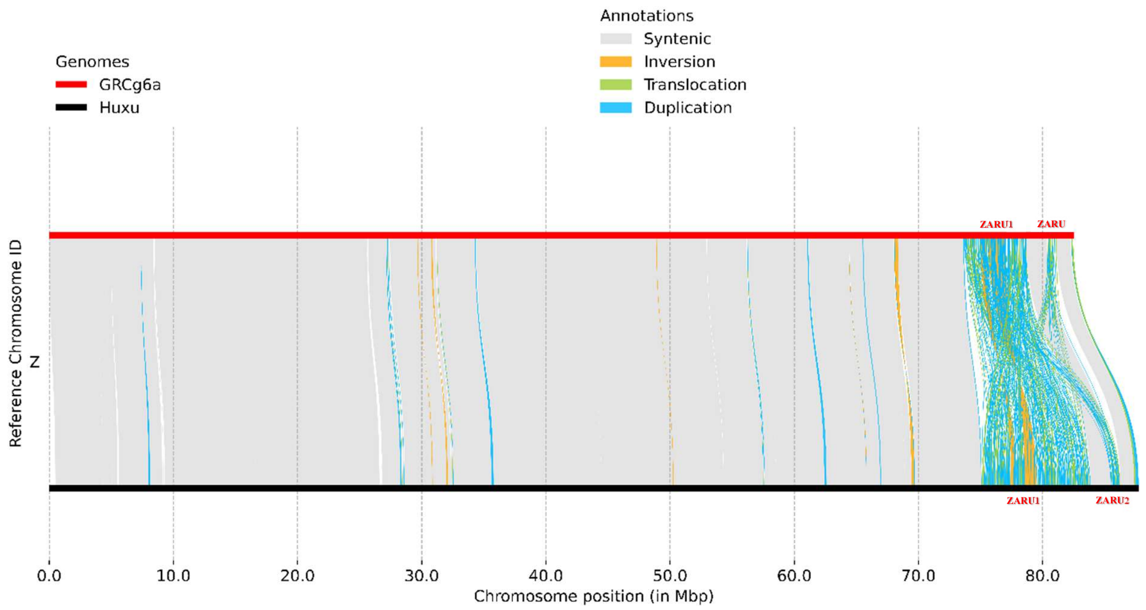


**Fig. S5: Organization of the chromosome *Z* in GRCg6a and Huxu genome assemblies:** The genomic organization of chromosome Z is elucidated through a visual depiction of the pairwise genome alignment between the GRCg6a and the Huxu genome. This alignment was performed using minimap2, syri, and plotsr. The figure highlights the ZARU1, Int, and ZARU2 regions.

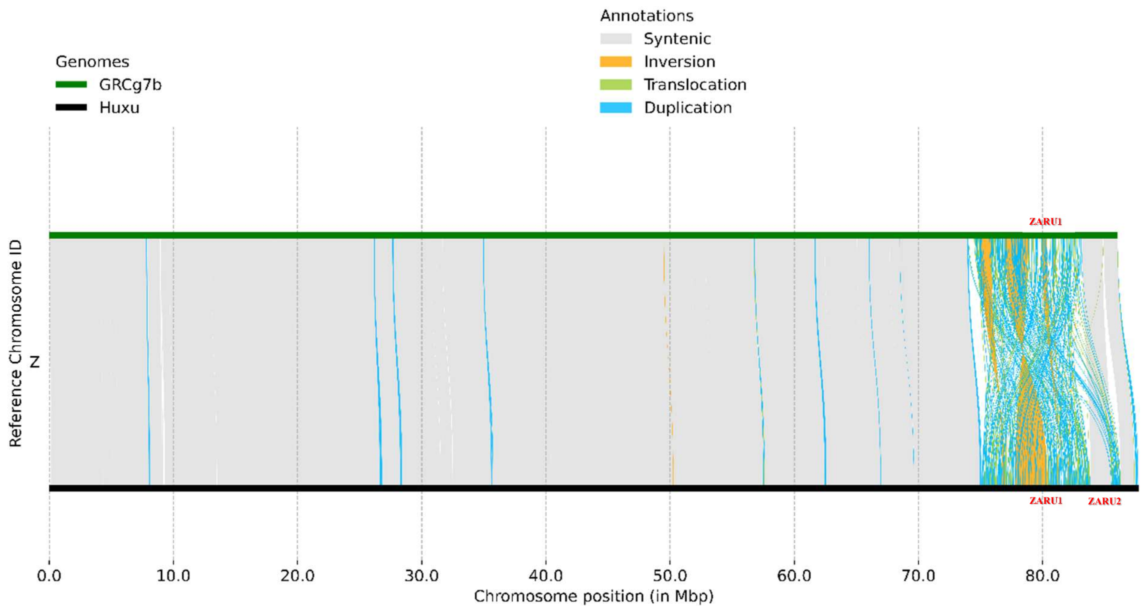


**Fig. S6: Organization of the chromosome *Z* in GRCg7b and Huxu genome assemblies:** The genomic organization of chromosome Z is elucidated through a visual depiction of the pairwise genome alignment between the GRCg7b and the Huxu genome. This alignment was performed using minimap2, syri, and plotsr. The figure highlights the ZARU1, Int, and ZARU2 regions.

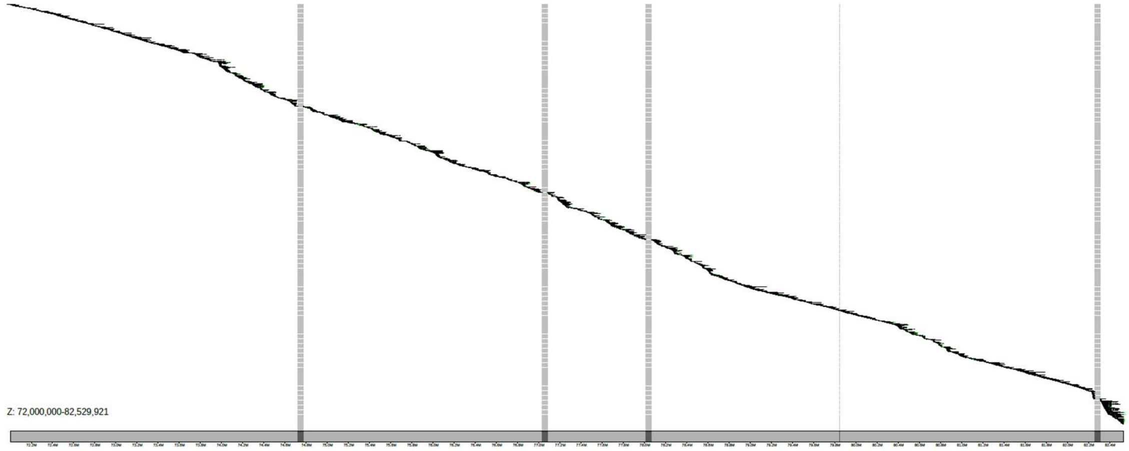


**Fig. S7: Evaluation of GRCg6a genome assembly using ONT long reads:** Klumpy scan alignment plot of ONT long-reads (SRR15421342, SRR15421343, SRR15421344, SRR15421345, SRR15421346, SRR13494713, and SRR13494714) aligned to the GRCg6a genome assembly using minimap2.

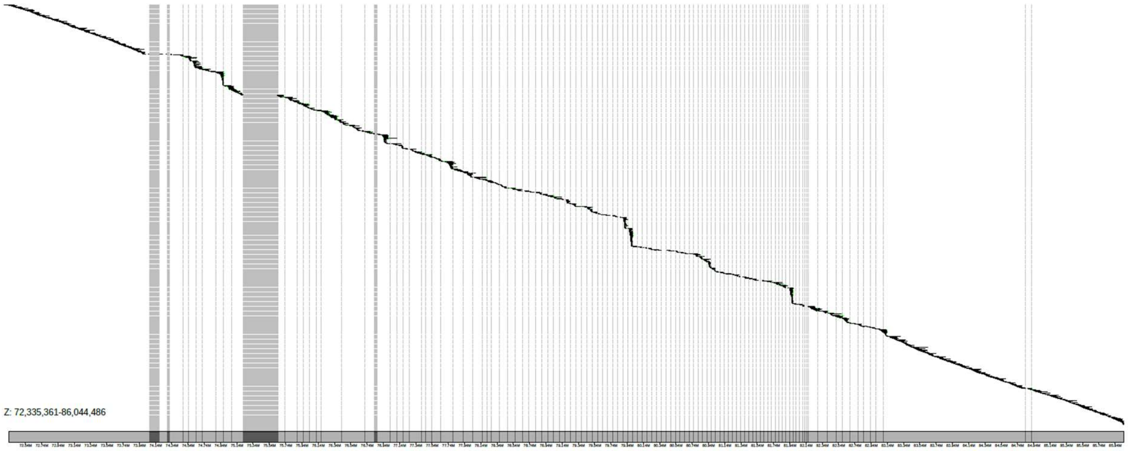


**Fig. S8: Evaluation of GRCg7b genome assembly using ONT long reads:** Klumpy scan alignment plot of ONT long-reads (SRR15421342, SRR15421343, SRR15421344, SRR15421345, SRR15421346, SRR13494713, and SRR13494714) aligned to the GRCg7b genome assembly using minimap2.

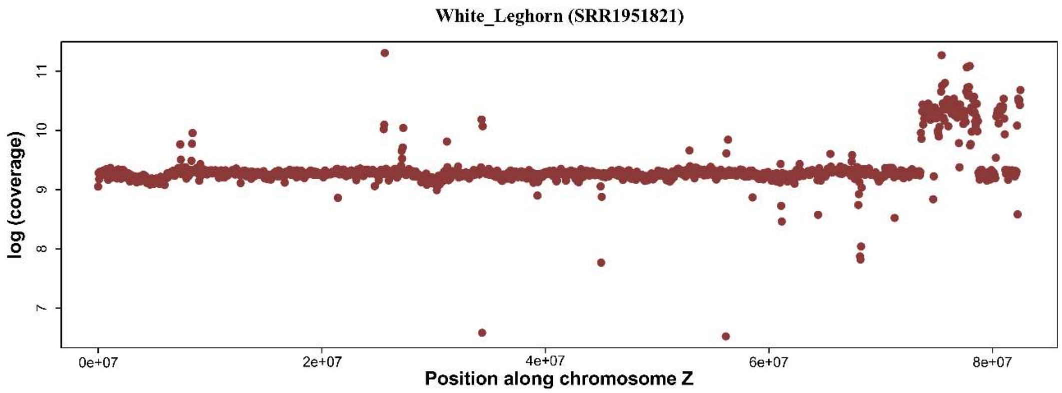


**Fig. S9:** The figure depicts the short-read coverage of the entire chromosome Z in a 10 Kb non-overlapping window for a white leghorn individual (SRR1951821) aligned with the GRCg6a genome assembly. The X-axis indicates the genomic position along chromosome Z, while the Y-axis represents the base log coverage of read counts in 10 Kb non-overlapping windows.

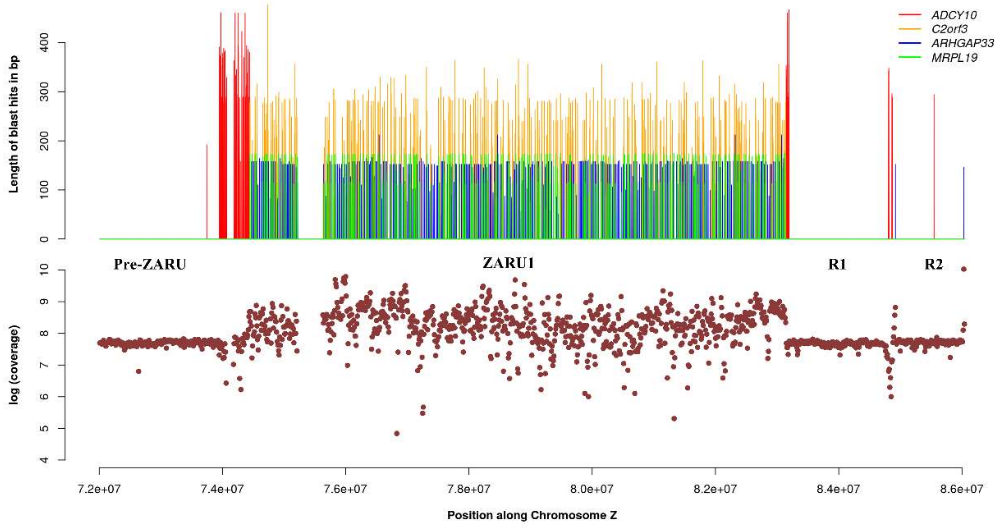


**Fig. S10:** A magnified perspective of a ~14 Mb segment (coordinates 72000000-86044486) located on the q-arm of Chicken chromosome Z in the GRCg7b genome assembly. The first panel displays the Z amplicon repeat units (ZARU), housing multiple copies of four genes: *ADCY10*, *C2orf3*, *ARHGAP33*, and *MRPL19*. In the second panel, the read coverage of a white leghorn individual (SRR1951821) aligned with the GRCg7b genome assembly within a 10 Kb window of the same region is depicted, indicating significant coverage attributed to repeat regions. The X-axis denotes the genomic position (Z:72000000-86044486) along chromosome Z. At the same time, the Y-axis represents the coverage of blast hit length in 1 Kb non-overlapping windows in the first panel, and the base log coverage of read counts in 10 Kb non-overlapping windows in the second panel.

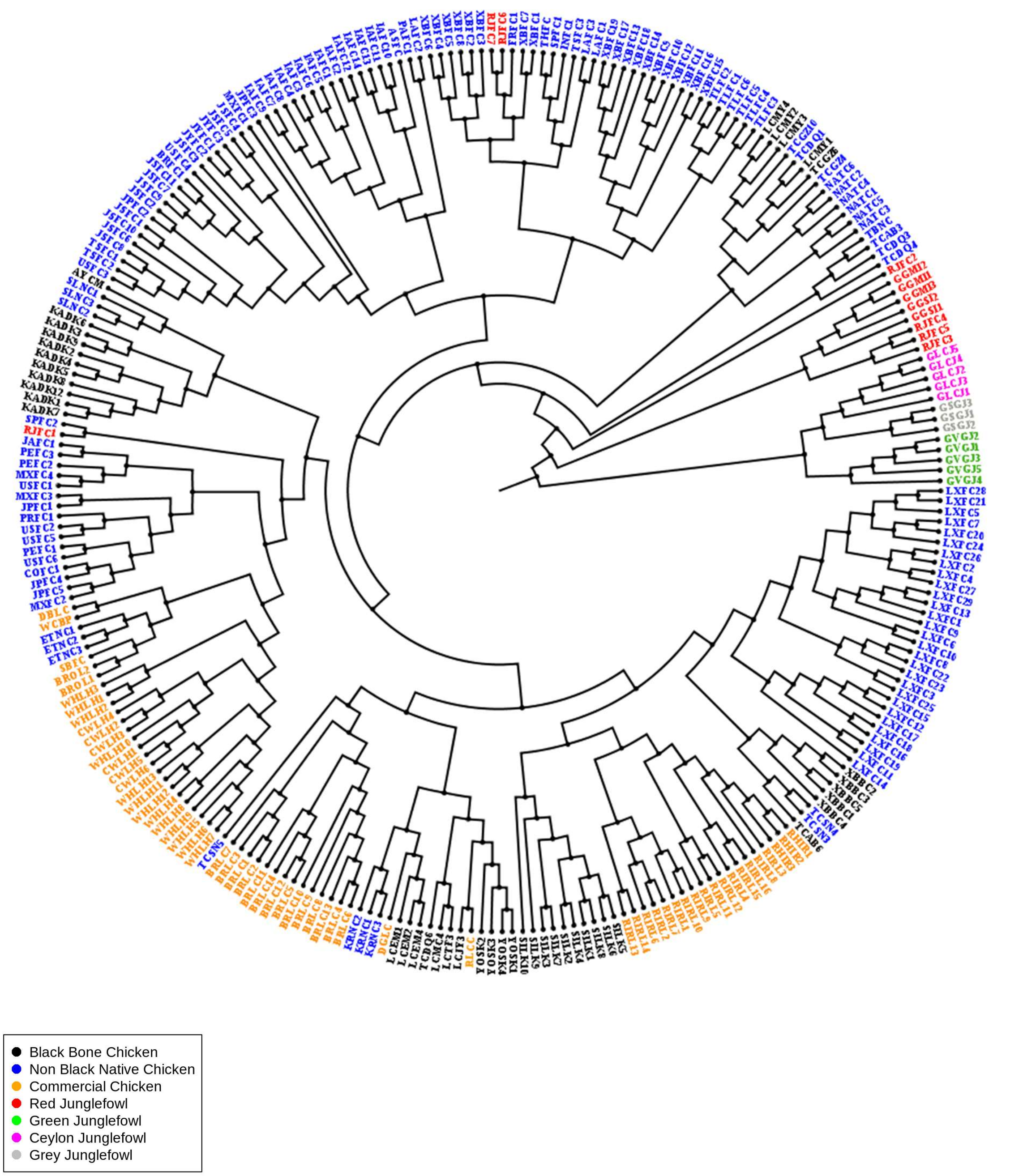


**Fig. S11:** Phylogenetic tree constructed using SNPhylo based on genome-wide SNPs (Single Nucleotide Polymorphisms), showcasing 270 individuals (42 Black bone chickens and 228 non-black chickens) and visualized using FigTree.

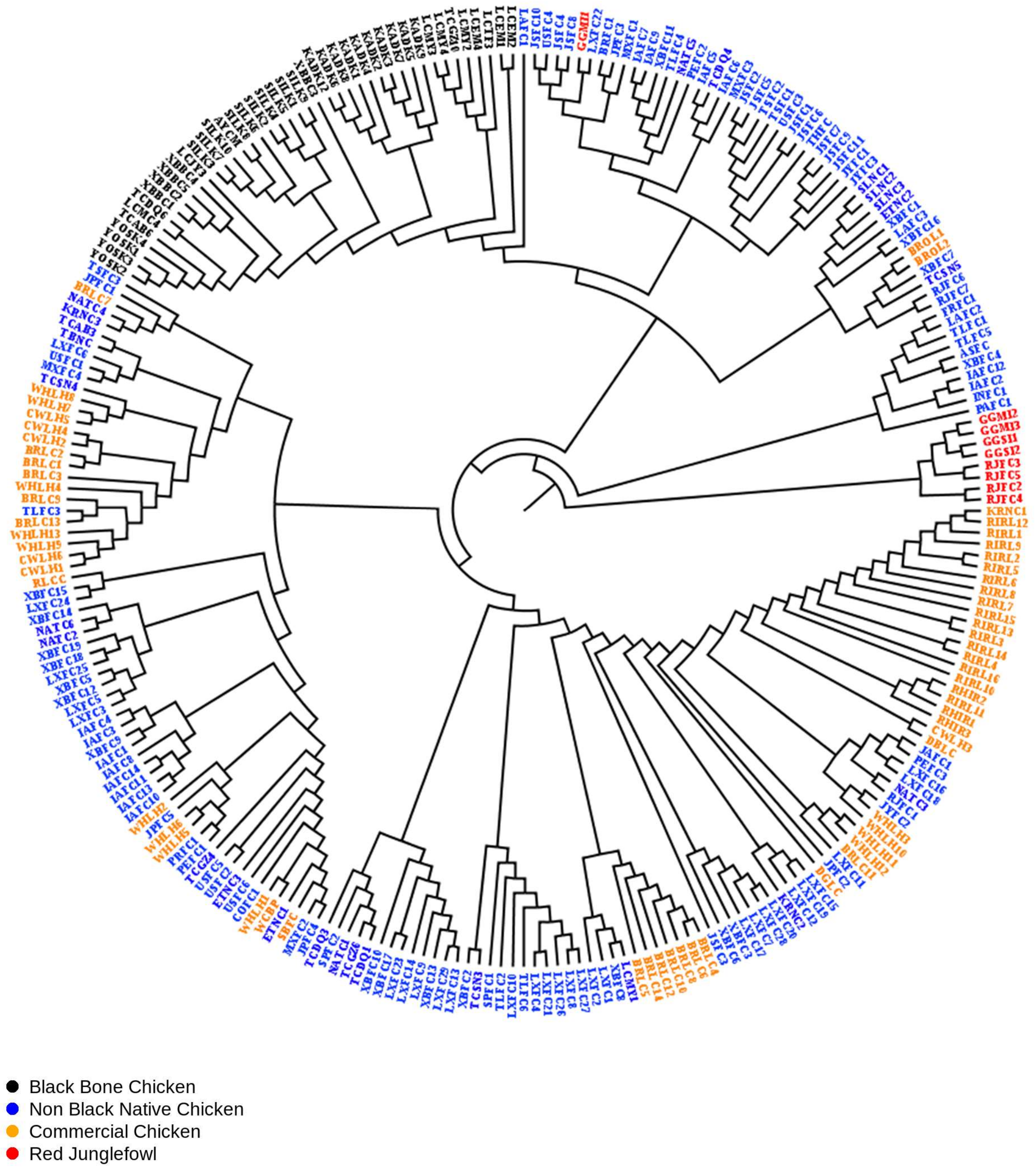


**Fig. S12:** Phylogenetic tree constructed using IQ-TREE2 based on SNPs (Single Nucleotide Polymorphisms) of *Fm* locus, showcasing 257 individuals (42 Black bone chickens and 215 non-black chickens) and visualized using FigTree.

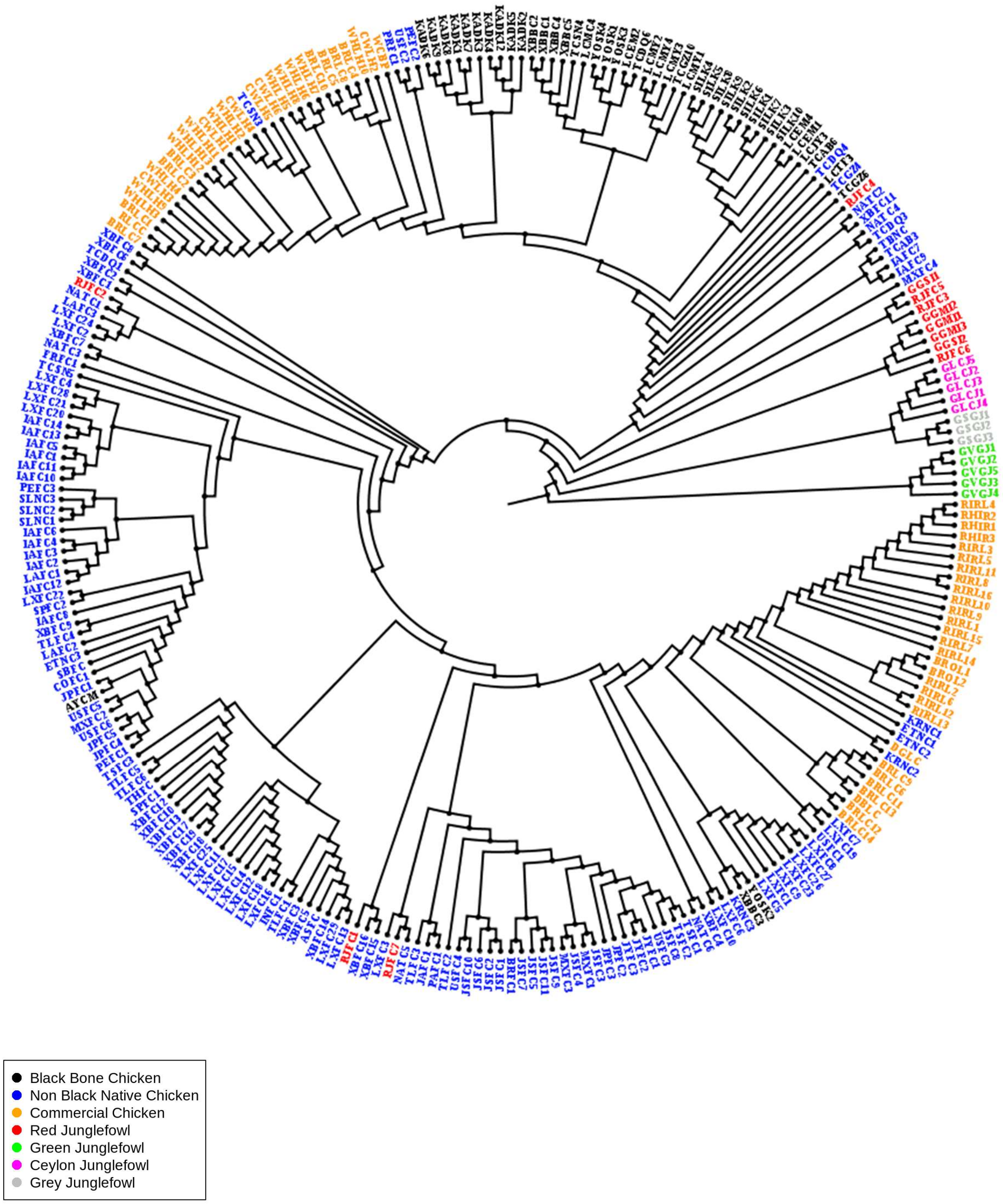


**Fig. S13:** Phylogenetic tree constructed using IQ-TREE2 based on SNPs (Single Nucleotide Polymorphisms) of the R1 region, showcasing 270 individuals (42 Black bone chickens and 228 non-black chickens) and visualized using FigTree.

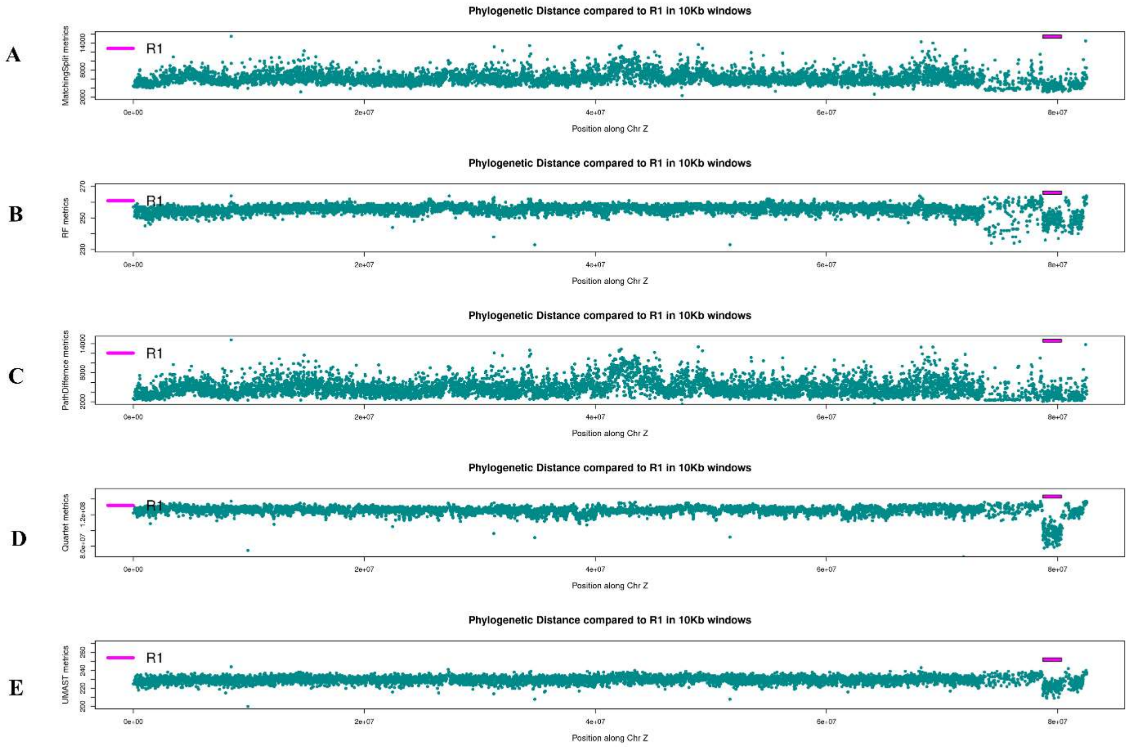


**Fig. S14:** phylogenetic distance metrics comparison between each of the 10Kb window trees of entire chromosome Z and R1 region phylogeny by using 5 metrics **(A)** MatchingSplit, **(B)** Robinson–Foulds, **(C)** PathDifference, **(D)** Quartet, and **(E)** UMAST in TreeCmp reveals that topologies of R1 exhibit the lowest phylogenetic distance. The pink rectangular box highlights the R1 region, while dark cyan-filled circles indicate the calculated distance metrics for each 10 Kb window along the Z chromosome.

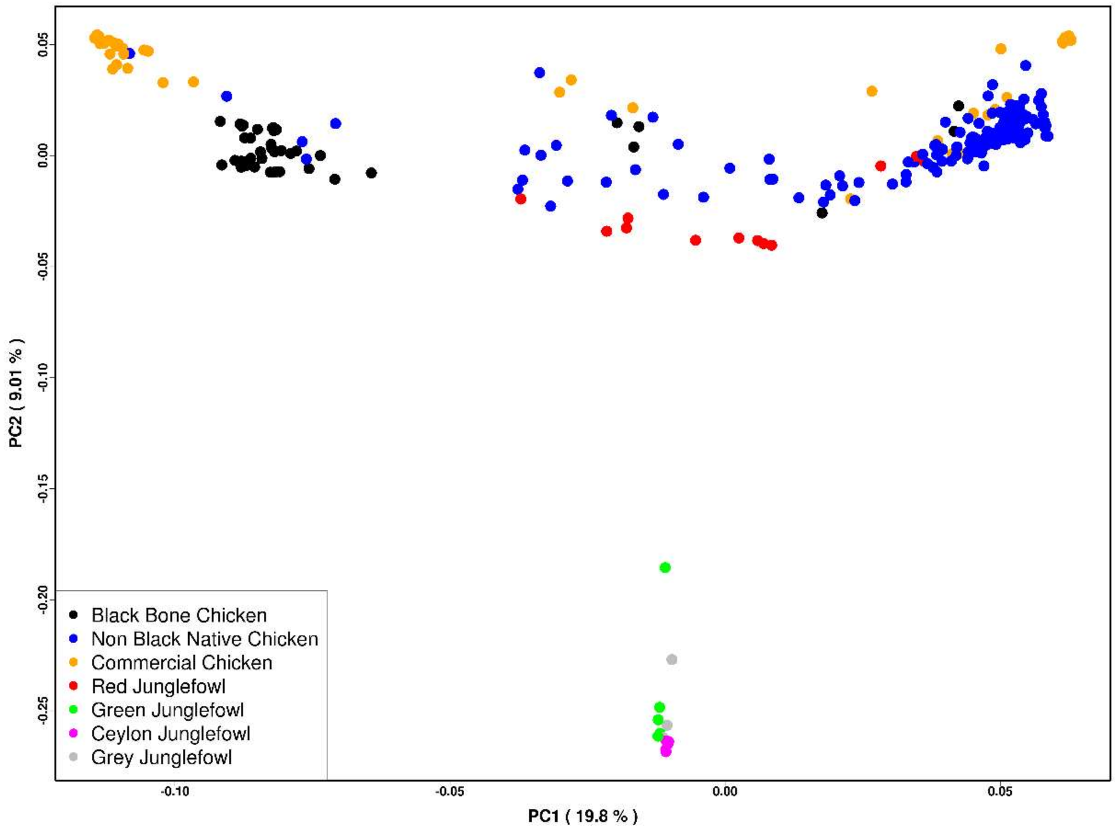


**Fig. S15:** Principal Component Analysis (PCA) plot illustrating the genetic relationships among 270 chicken individuals based on the R1 region. PC1 and PC2 account for 19.8% and 9.01% of the variance, respectively. Black-bone breeds are represented in black, while non-black-bone breeds, including wild junglefowl species, are indicated by various distinct colours.

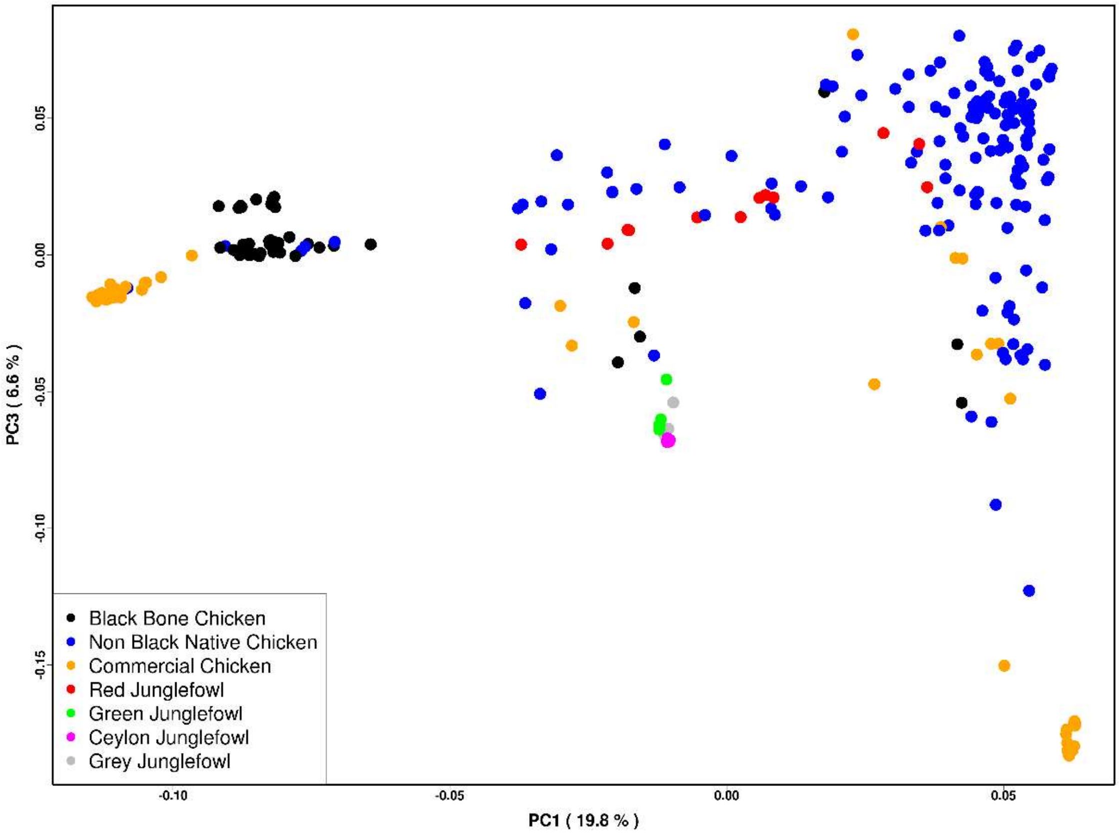


**Fig. S16:** Principal Component Analysis (PCA) plot illustrating the genetic relationships among 270 chicken individuals based on the R1 region. PC1 and PC3 account for 19.8% and 6.6% of the variance, respectively. Black-bone breeds are represented in black, while non-black-bone breeds, including wild junglefowl species, are indicated by various distinct colours.

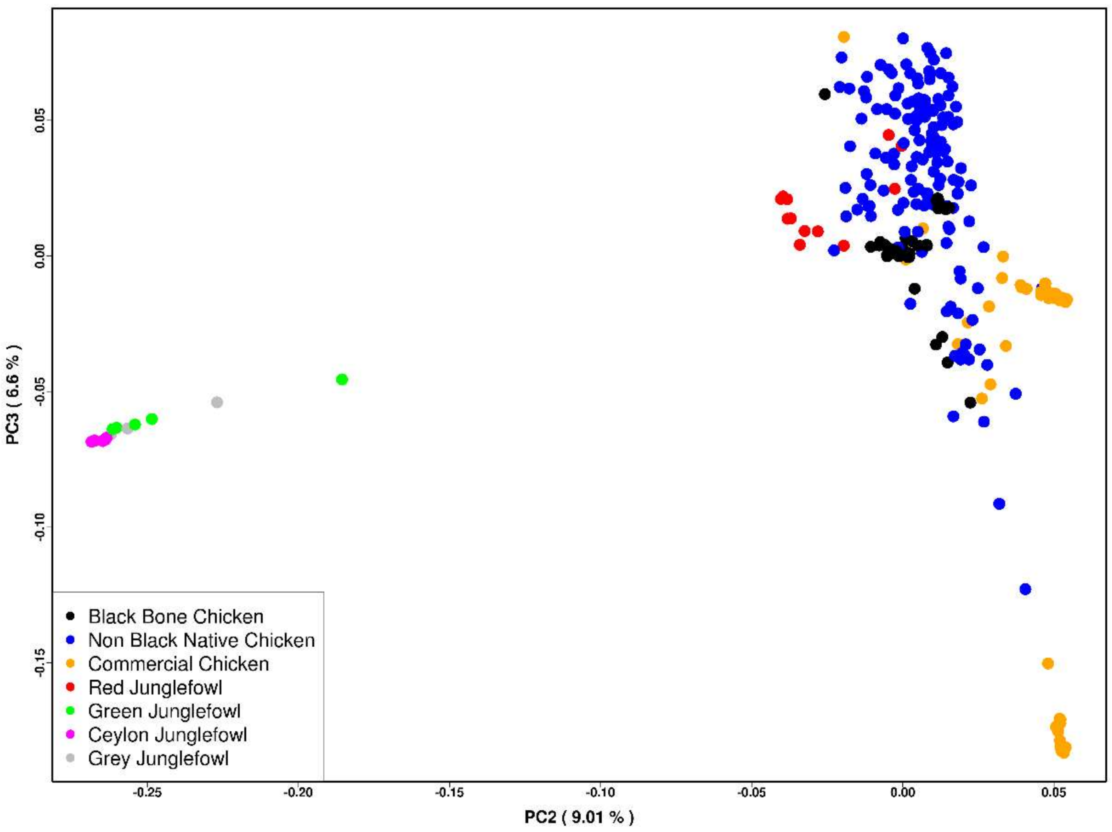


**Fig. S17:** Principal Component Analysis (PCA) plot illustrating the genetic relationships among 270 chicken individuals based on the R1 region. PC2 and PC3 account for 9.01% and 6.6% of the variance, respectively. Black-bone breeds are represented in black, while non-black-bone breeds, including wild junglefowl species, are indicated by various distinct colours.

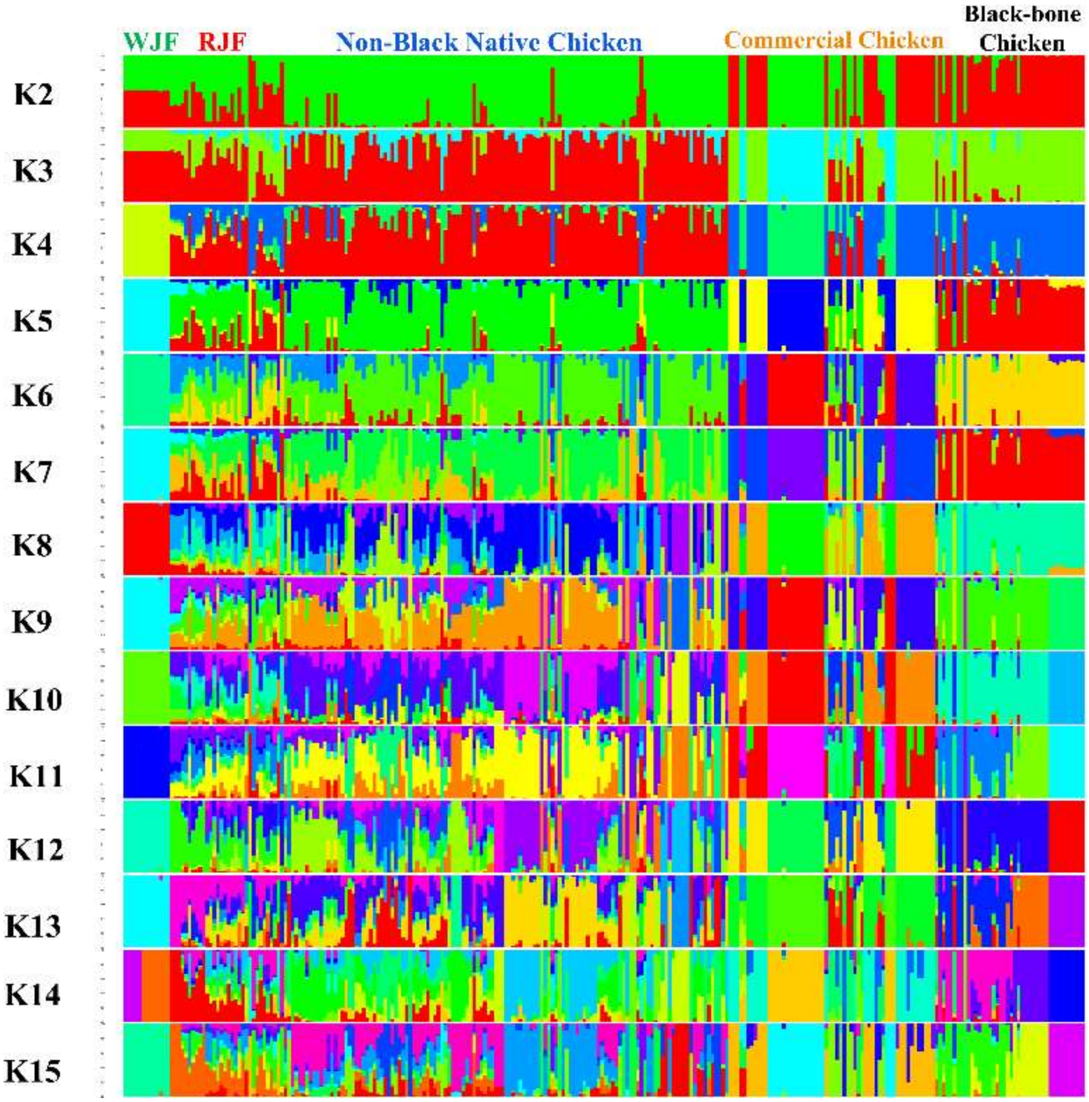


**Fig. S18:** Genetic Structure and Individual Ancestry in the R1 Region: This figure illustrates the genetic structure and individual ancestry analysis conducted on a dataset of 270 chickens from various breeds. The analysis was carried out using the NGSadmix software and encompassed a range of K values, from K=2 to K=15, revealing the genetic composition and clustering patterns of the studied individuals.

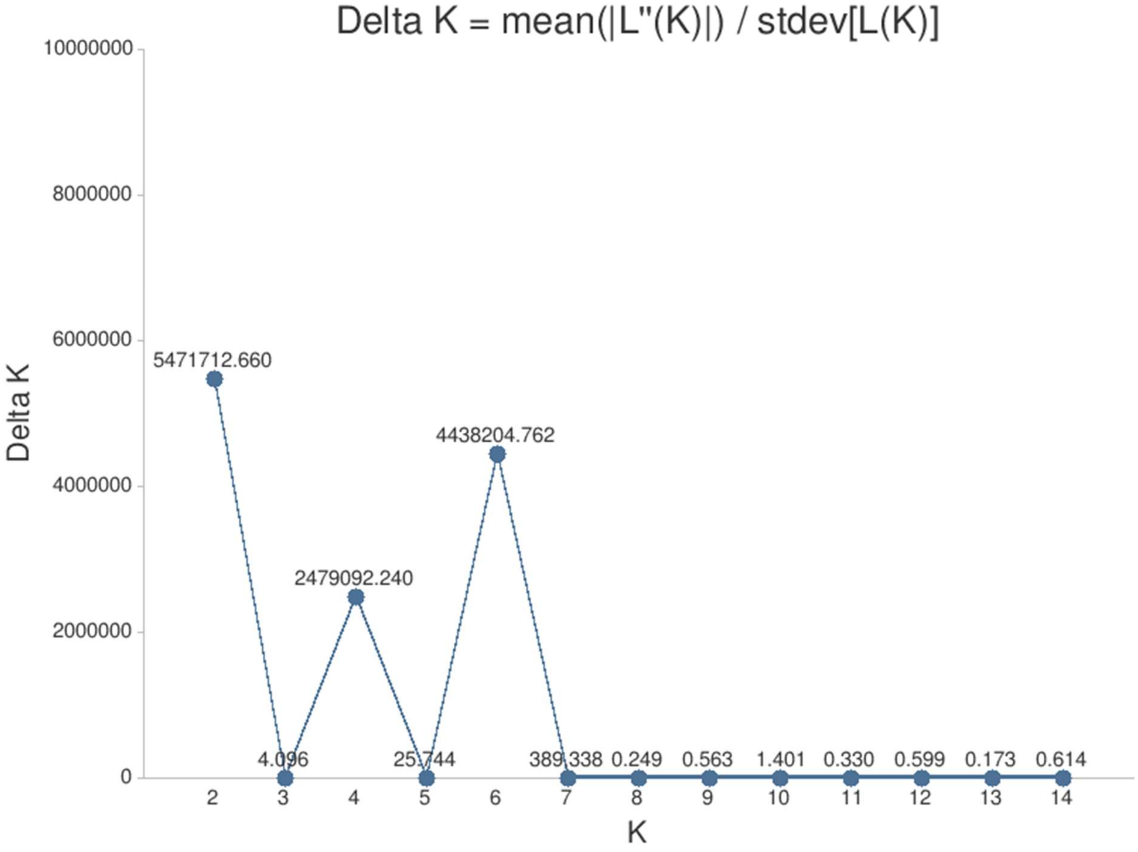


**Fig. S19:** A graphical representation displaying the Delta K values obtained from the CLUMPAK web server for the admixture analysis of the R1 region.

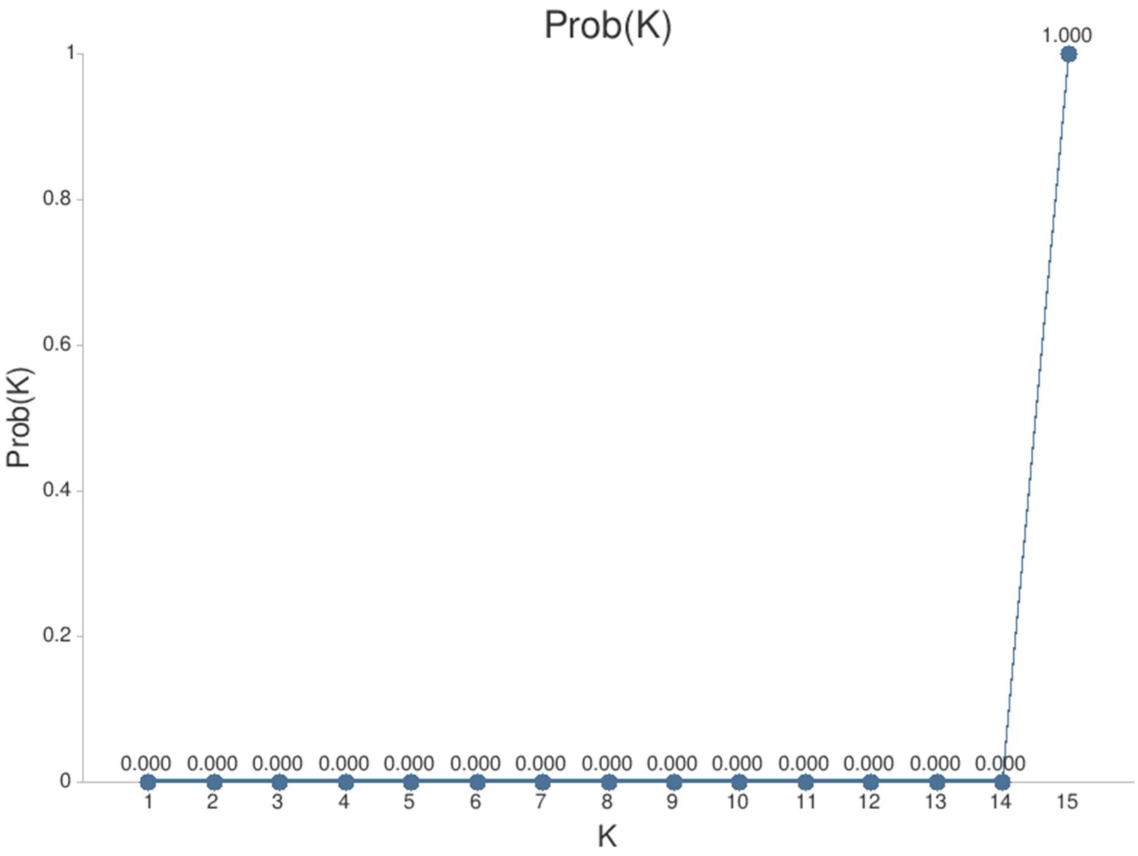


**Fig. S20**: Log probability of the data referred to as ln Pr(X|K) value plot obtained from the CLUMPAK web server for the admixture analysis of the R1 region.

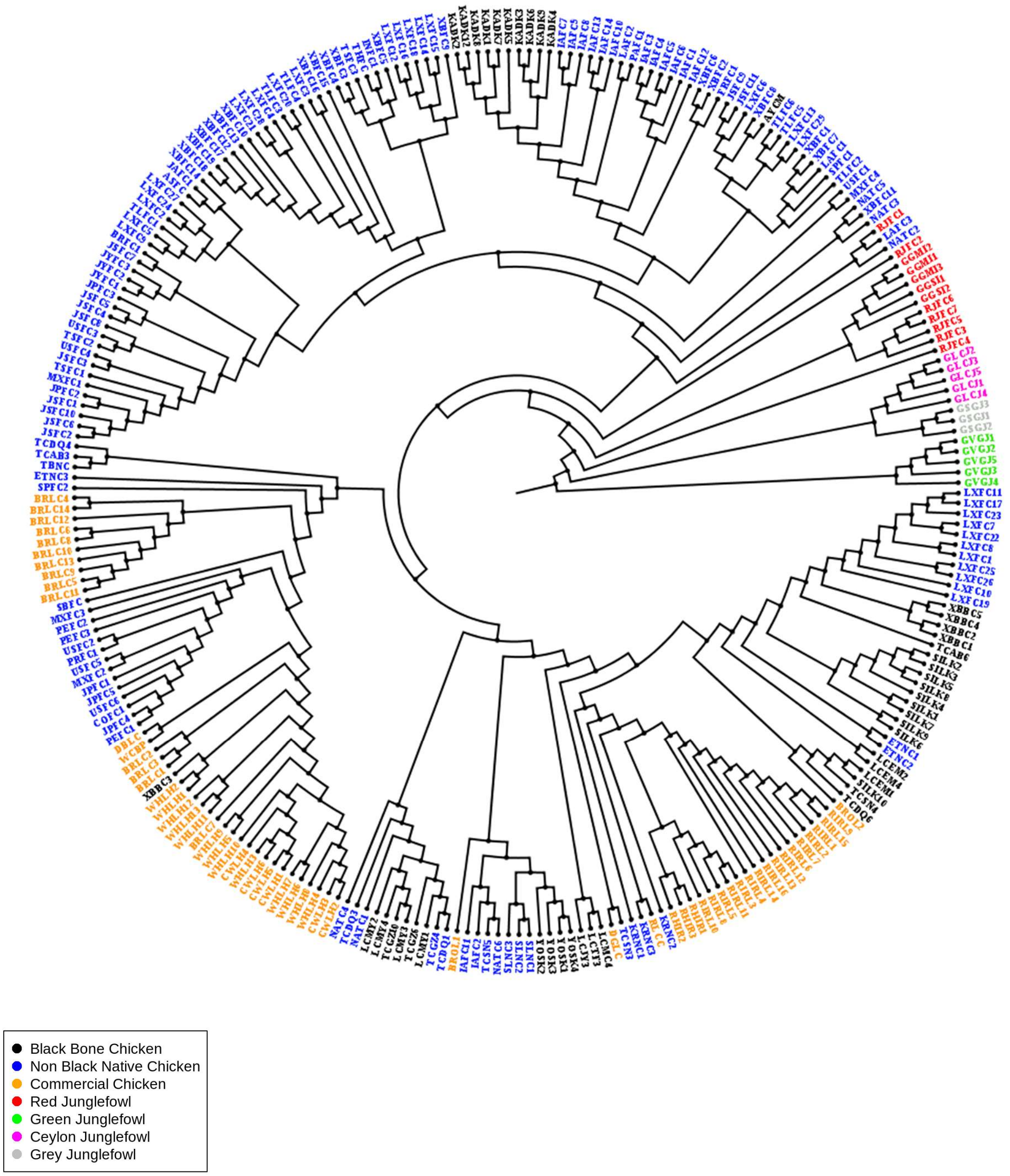


**Fig. S21:** Phylogenetic tree constructed using IQ-TREE2 based on SNPs (Single Nucleotide Polymorphisms) of the R2 region, showcasing 270 individuals (42 Black bone chickens and 228 non-black chickens) and visualized using FigTree.

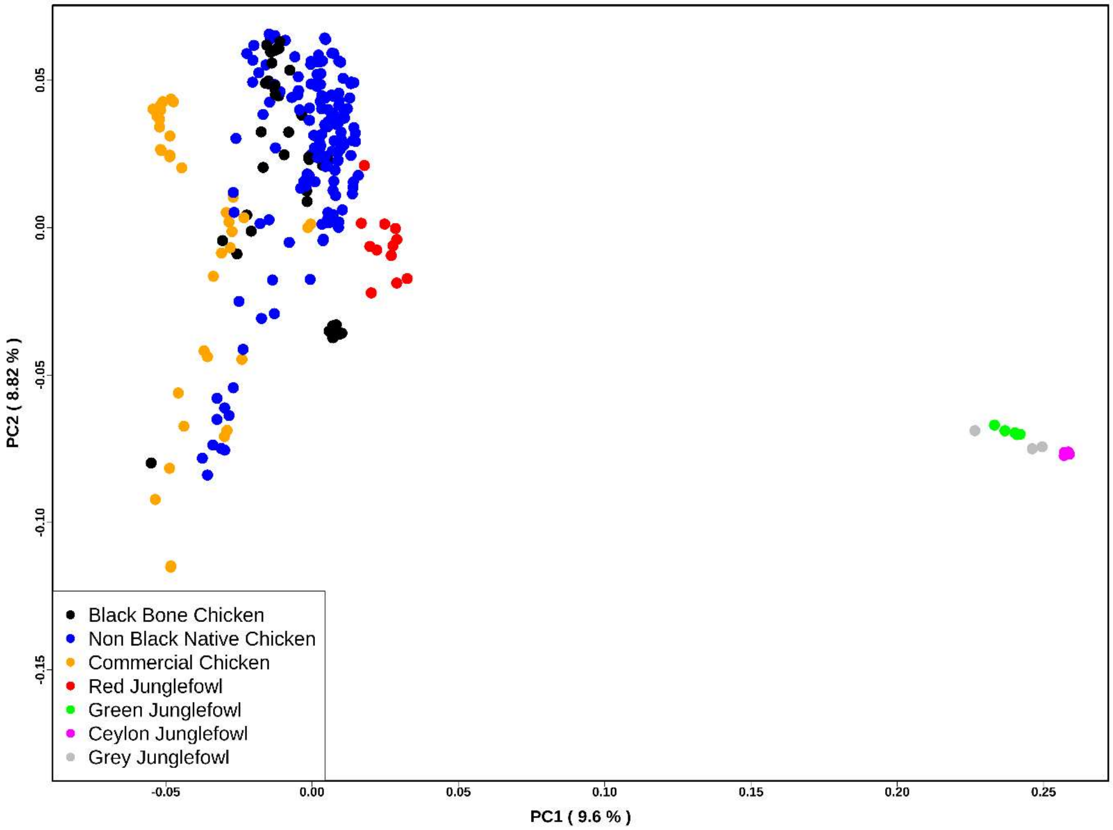


**Fig. S22:** Principal Component Analysis (PCA) plot illustrating the genetic relationships among 270 chicken individuals based on the R2 region. PC1 and PC2 account for 9.6% and 8.82% of the variance, respectively. Black-bone breeds are represented in black, while non-black-bone breeds, including wild junglefowl species, are indicated by various distinct colours.

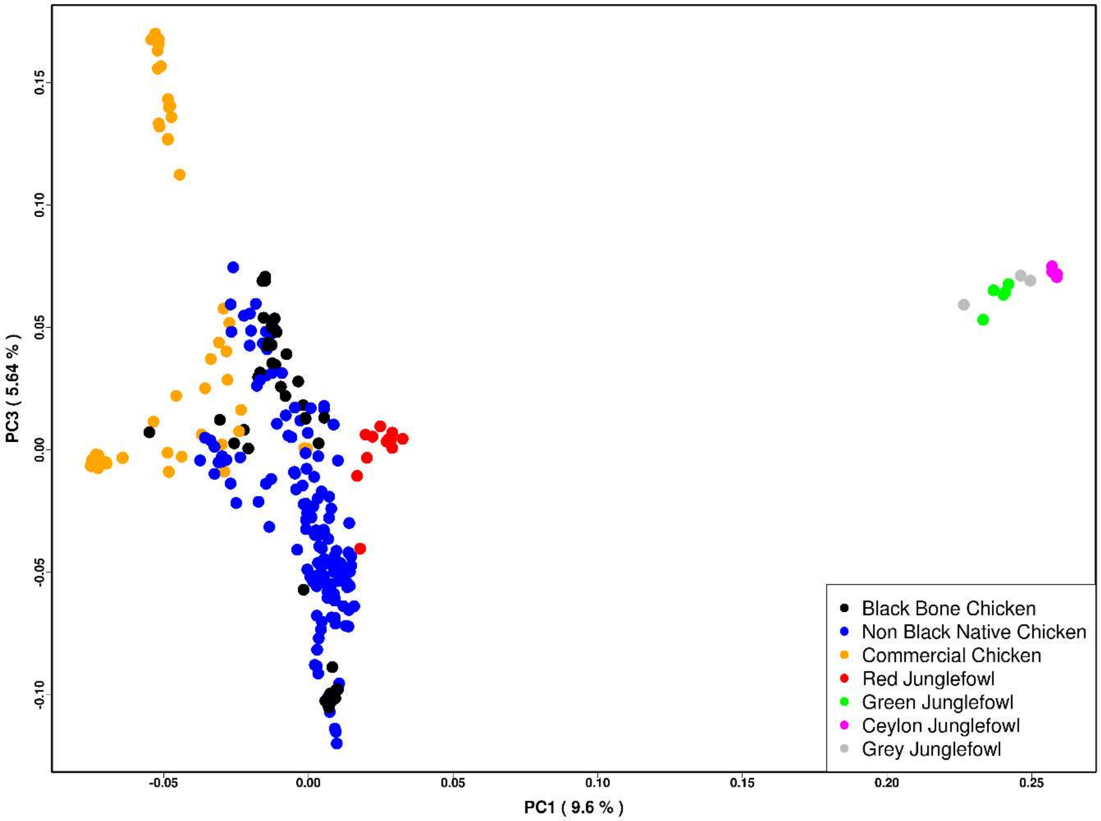


**Fig. S23:** Principal Component Analysis (PCA) plot illustrating the genetic relationships among 270 chicken individuals based on the R2 region. PC1 and PC3 account for 9.6% and 6.64% of the variance, respectively. Black-bone breeds are represented in black, while non-black-bone breeds, including wild junglefowl species, are indicated by various distinct colours.

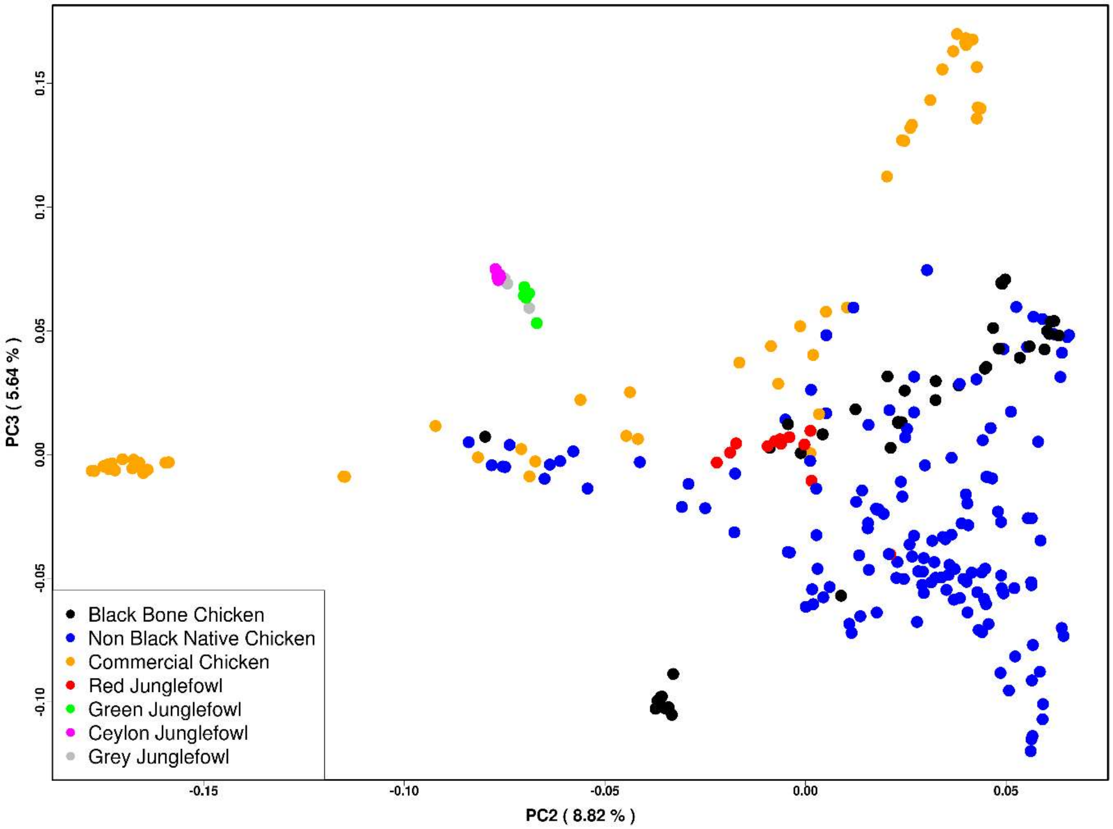


**Fig. S24:** Principal Component Analysis (PCA) plot illustrating the genetic relationships among 270 chicken individuals based on the R2 region. PC2 and PC3 account for 8.82% and 6.64% of the variance, respectively. Black-bone breeds are represented in black, while non-black-bone breeds, including wild junglefowl species, are indicated by various distinct colours.

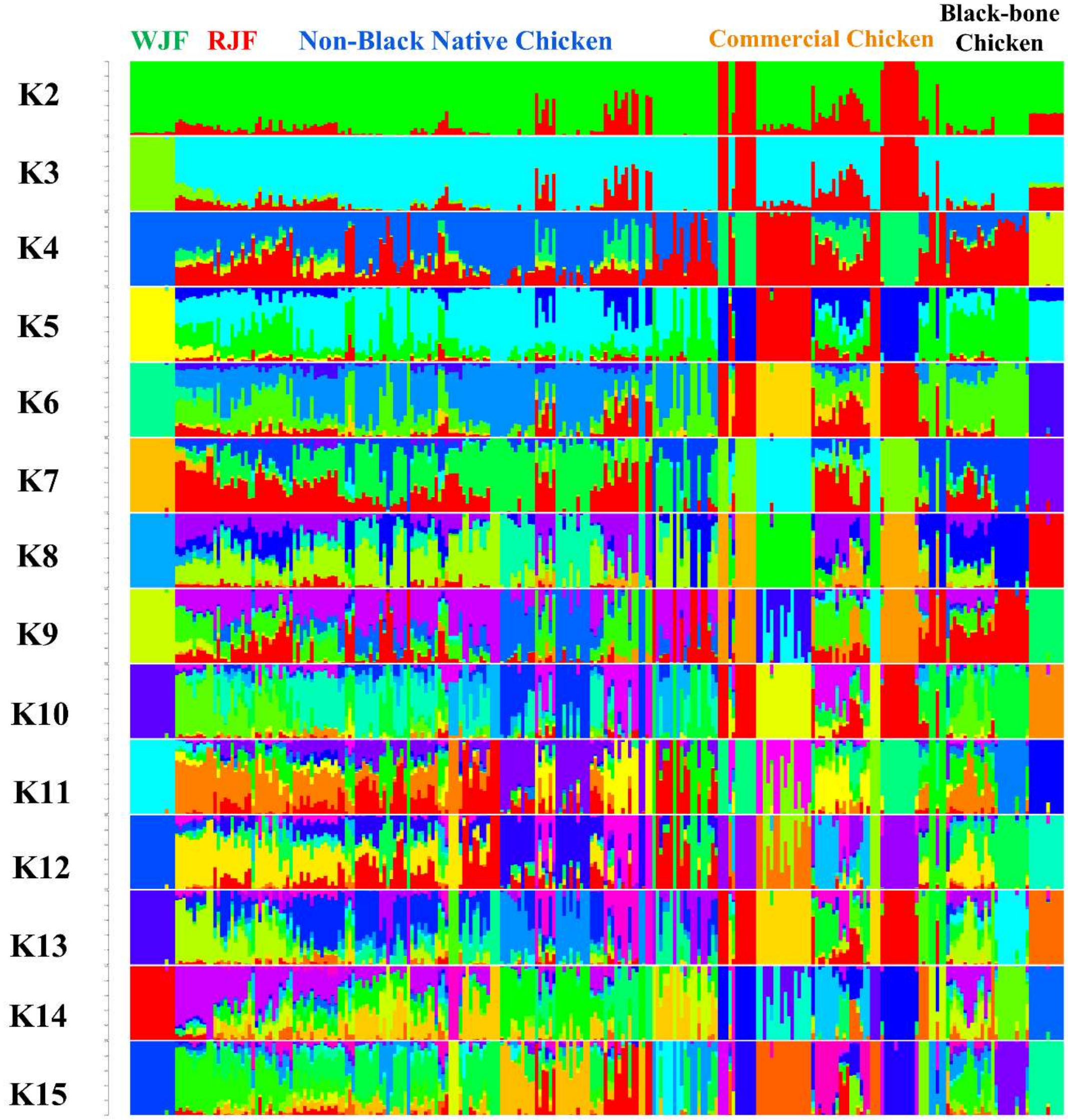


**Fig. S25: Genetic Structure and Individual Ancestry in the R2 Region:** This figure illustrates the genetic structure and individual ancestry analysis conducted on a dataset of 270 chickens from various breeds. The analysis was carried out using the NGSadmix software and encompassed a range of K values, from K=2 to K=15, revealing the genetic composition and clustering patterns of the studied individuals.

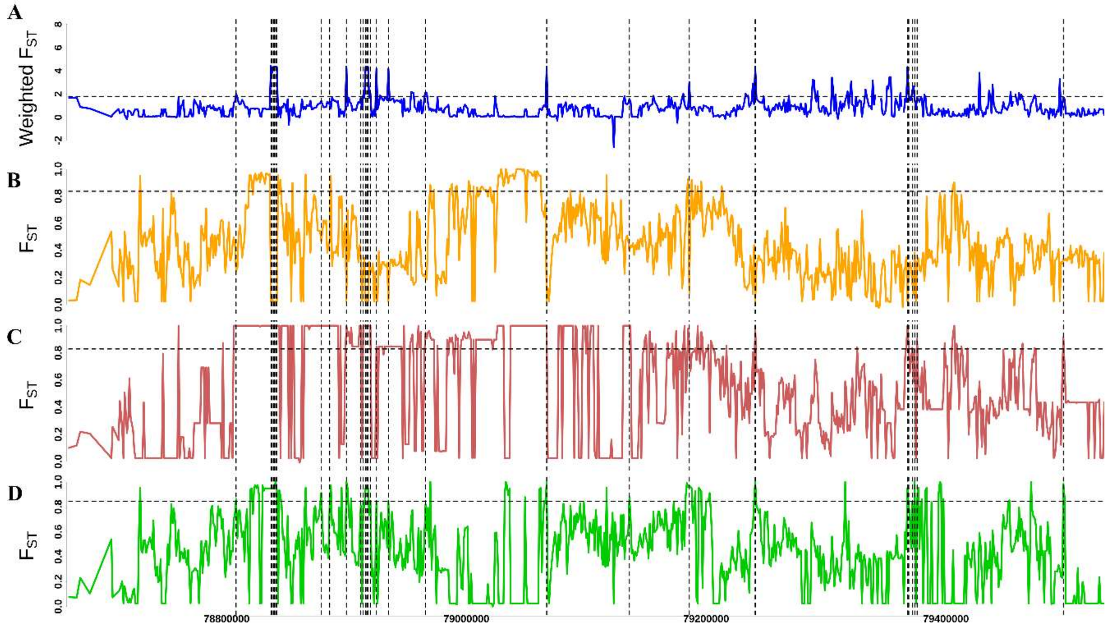


**Fig S26: Black bone chicken specific signature of selection in R1: (A)** Comparative analysis of Weighted $F_{ST}$ (KADK vs. WHLH + SILK vs. WHLH/KADK vs. SILK) between black bone chicken (KADK and SILK) and non-black bone chicken (WHLH). **(B)** $F_{ST}$ comparison between KADK and SILK. **(C)** $F_{ST}$ comparison between KADK and WHLH. **(D)** $F_{ST}$ comparison between SILK and WHLH. A horizontal black dotted line represents the 90th percentile threshold for $F_{ST}$ comparisons. while black dotted vertical lines represent the targeted region.

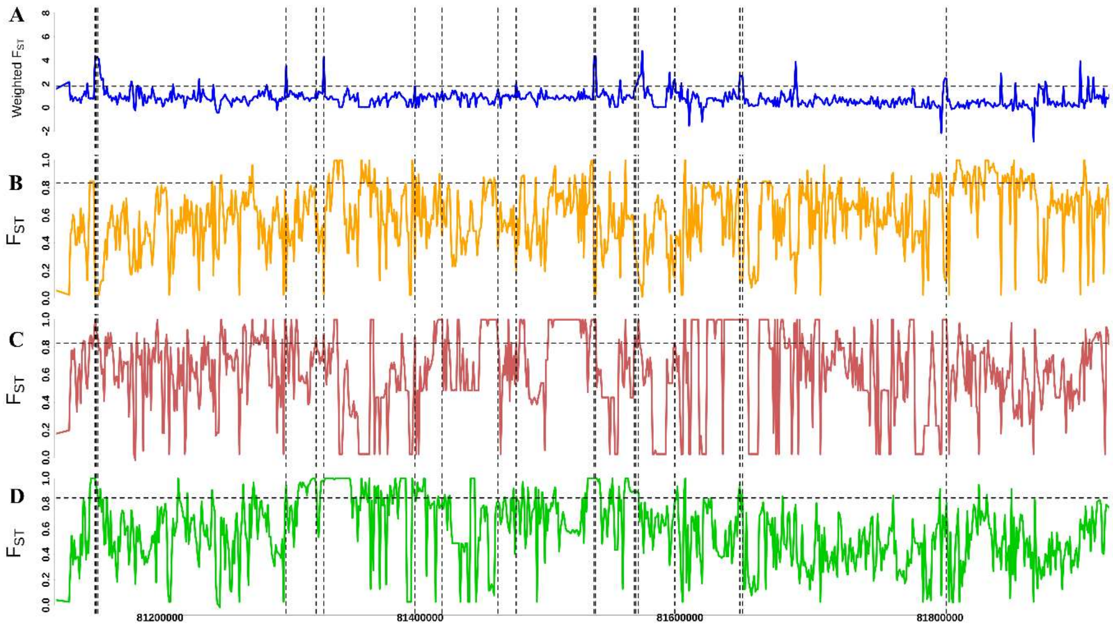


**Fig S27: Black bone chicken specific signature of selection in R2: (A)** Comparative analysis of Weighted $F_{ST}$ (KADK vs. WHLH + SILK vs. WHLH/KADK vs. SILK) between black bone chicken (KADK and SILK) and non-black bone chicken (WHLH). **(B)** $F_{ST}$ comparison between KADK and SILK. **(C)** $F_{ST}$ comparison between KADK and WHLH. **(D)** $F_{ST}$ comparison between SILK and WHLH. A horizontal black dotted line represents the 90th percentile threshold for $F_{ST}$ comparisons. while black dotted vertical lines represent the targeted region.

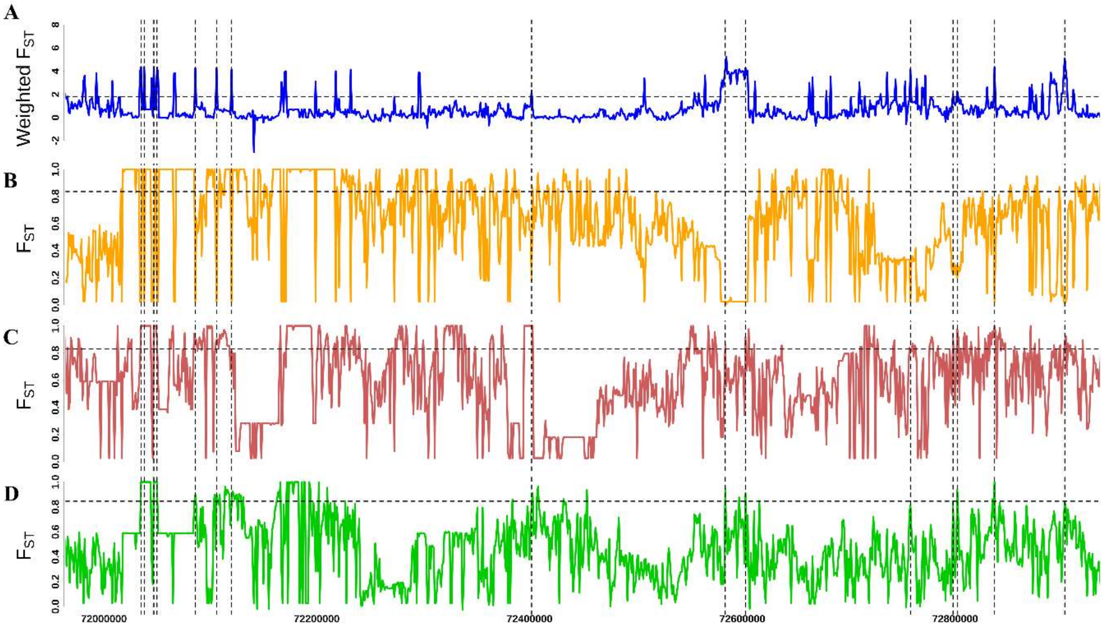


**Fig S28: Black bone chicken specific signature of selection in the upstream region of ZARU: (A)** Comparative analysis of Weighted $F_{ST}$ (KADK vs. WHLH + SILK vs. WHLH/KADK vs. SILK) between black bone chicken (KADK and SILK) and non-black bone chicken (WHLH). **(B)** $F_{ST}$ comparison between KADK and SILK. **(C)** $F_{ST}$ comparison between KADK and WHLH. **(D)** $F_{ST}$ comparison between SILK and WHLH. A horizontal black dotted line represents the 90th percentile threshold for $F_{ST}$ comparisons. while black dotted vertical lines represent the targeted region.

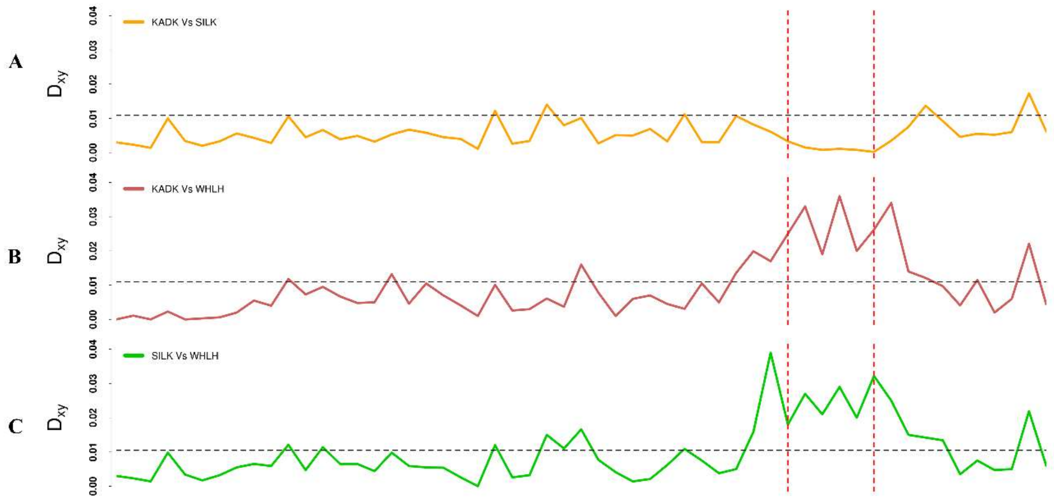


**Fig. S29:** Comparison of examined pairwise $D_{xy}$ between **(A)** BBC vs. BBC (KADK vs. SILK) and **(B and C)** BBC vs. non-BBC (KADK vs. WHLH, and SILK vs. WHLH) within the in the last intron of the *MTAP* gene located in R1. Dotted vertical red lines represent the boundaries of the targeted region.

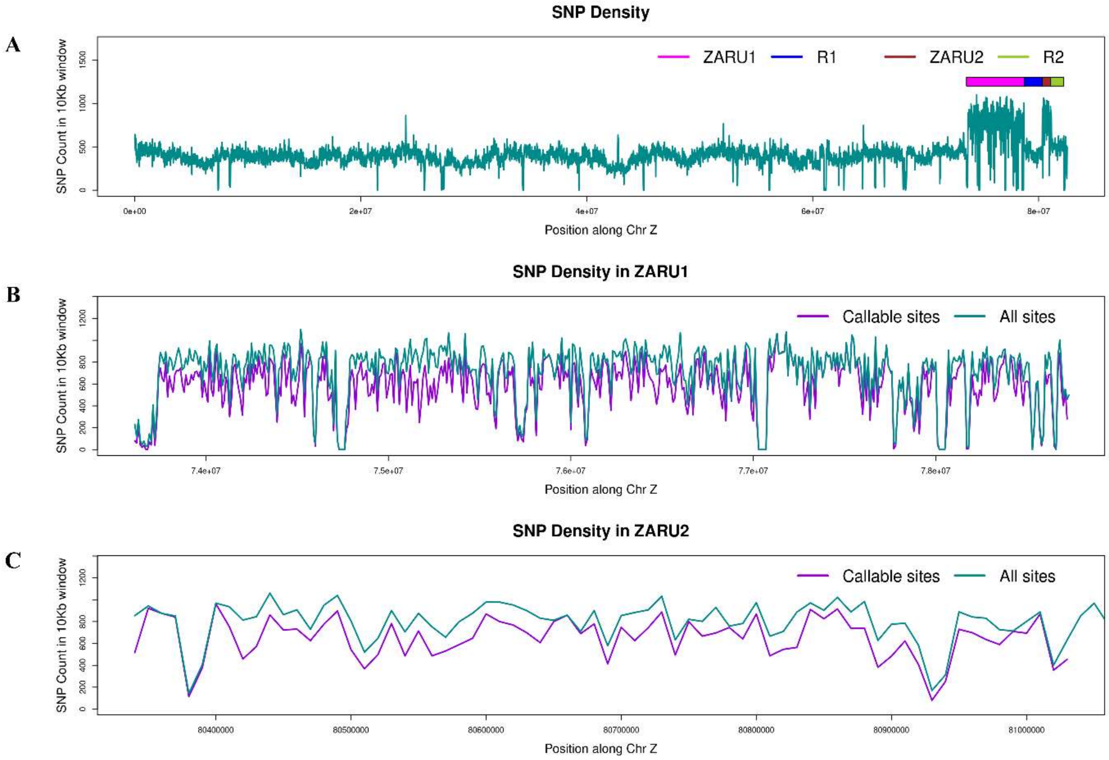


**Fig S30:** SNP density across chromosome Z**,** (**A**)The dark cyan solid line illustrates SNP density in 10 Kb non-overlapping windows across chromosome Z. Rectangular coloured boxes highlight specific regions, with dark violet representing the SNP density for callable sites within the same 10 Kb non-overlapping windows. (**B**) and (**C**) represents the SNP density in ZARU1 and ZARU2 regions, respectively.

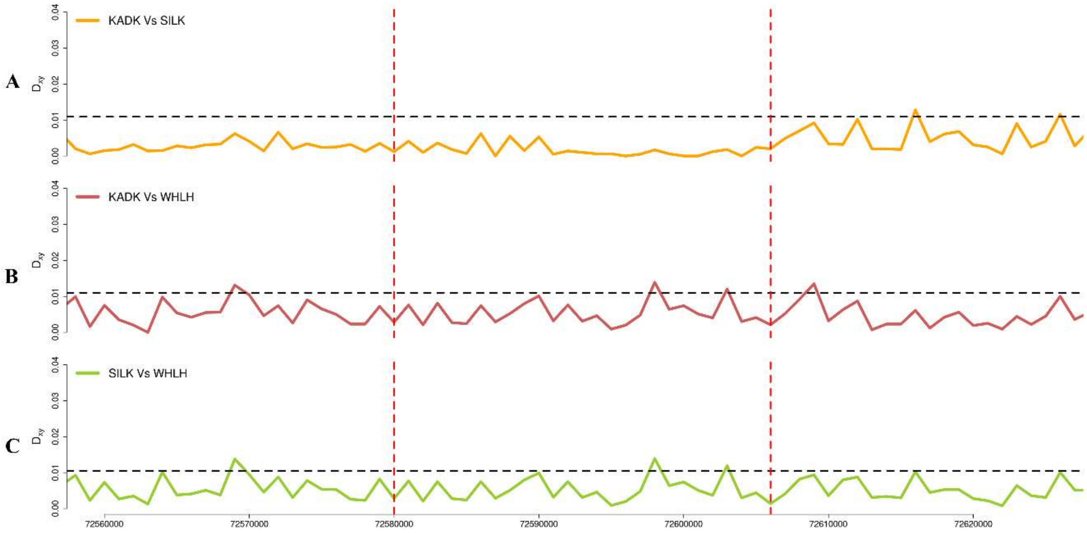


**Fig S31:** Comparison of examined pairwise $D_{xy}$ between **(A)** BBC vs. BBC (KADK vs. SILK) and **(B and C)** BBC vs. non-BBC (KADK vs. WHLH, and SILK vs. WHLH) within the 26 kb long intergenic region upstream to the ZARU. Dotted vertical red lines represent the boundaries of the targeted region.

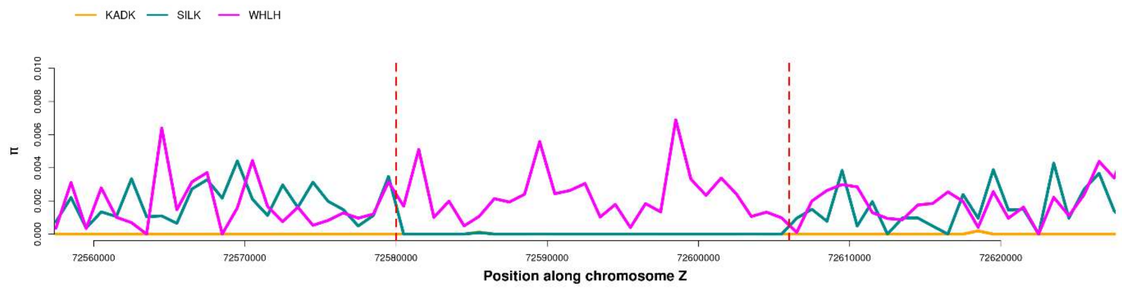


**Fig S32:** Comparison of examined nucleotide diversity (π) between BBC (KADK and SILK) and non-BBC (WHLH) within the 26 kb long intergenic region upstream to the ZARU. Dotted vertical red lines represent the boundaries of the targeted region.

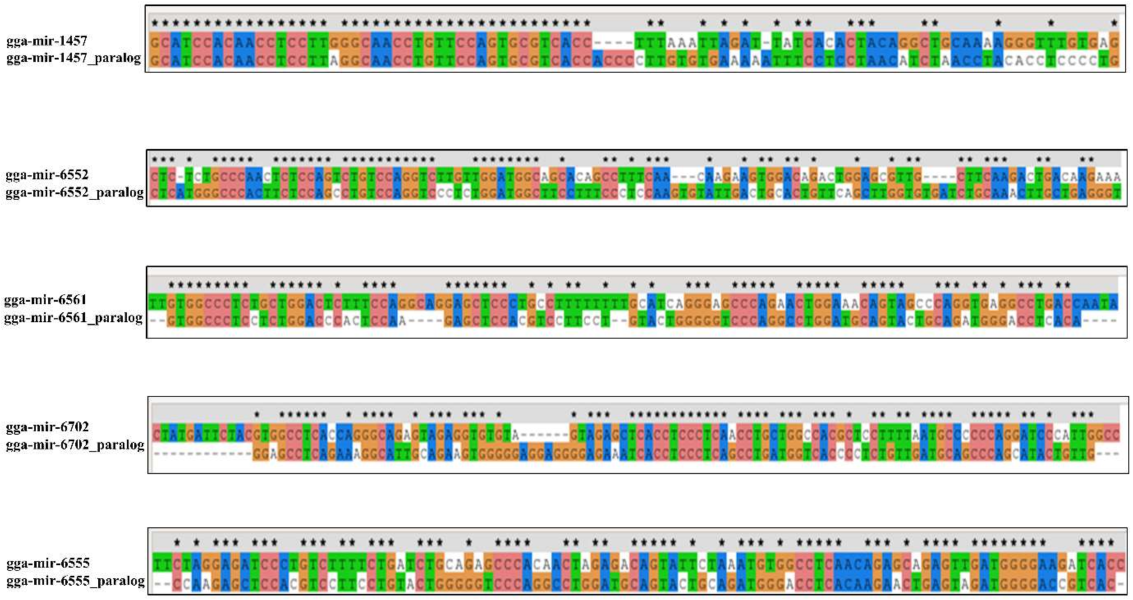


**Fig S33:** Sequence alignment of 5 microRNA paralogs identified in the 26 Kb long putatively selected region upstream to the Z amplicon repeat units.

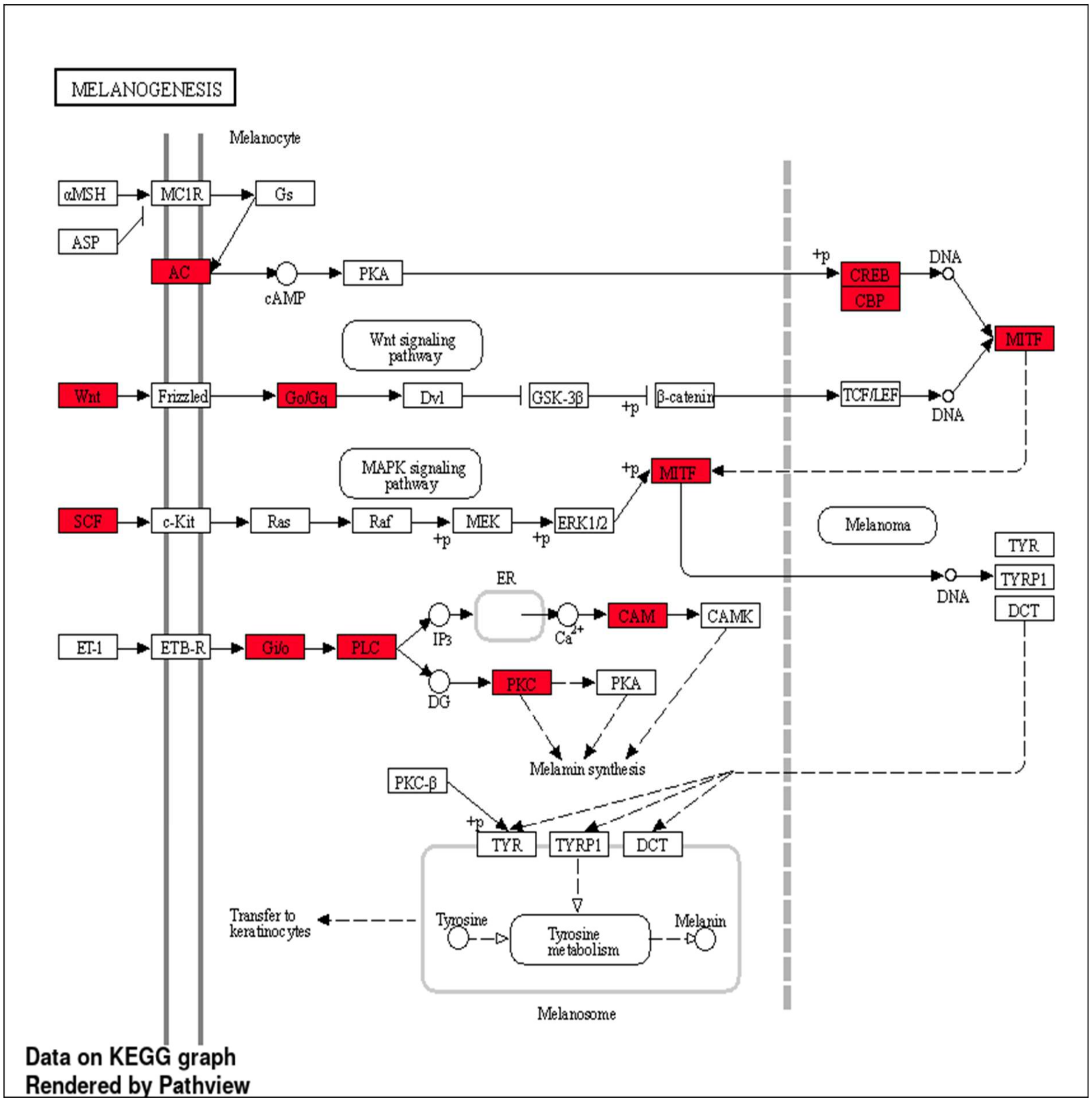


**Fig S34:** A diagram depicting the Melanogenesis pathway sourced from the KEGG pathway database, illustrating the relationship between the target genes regulated by the gga-mir-1457 paralog.

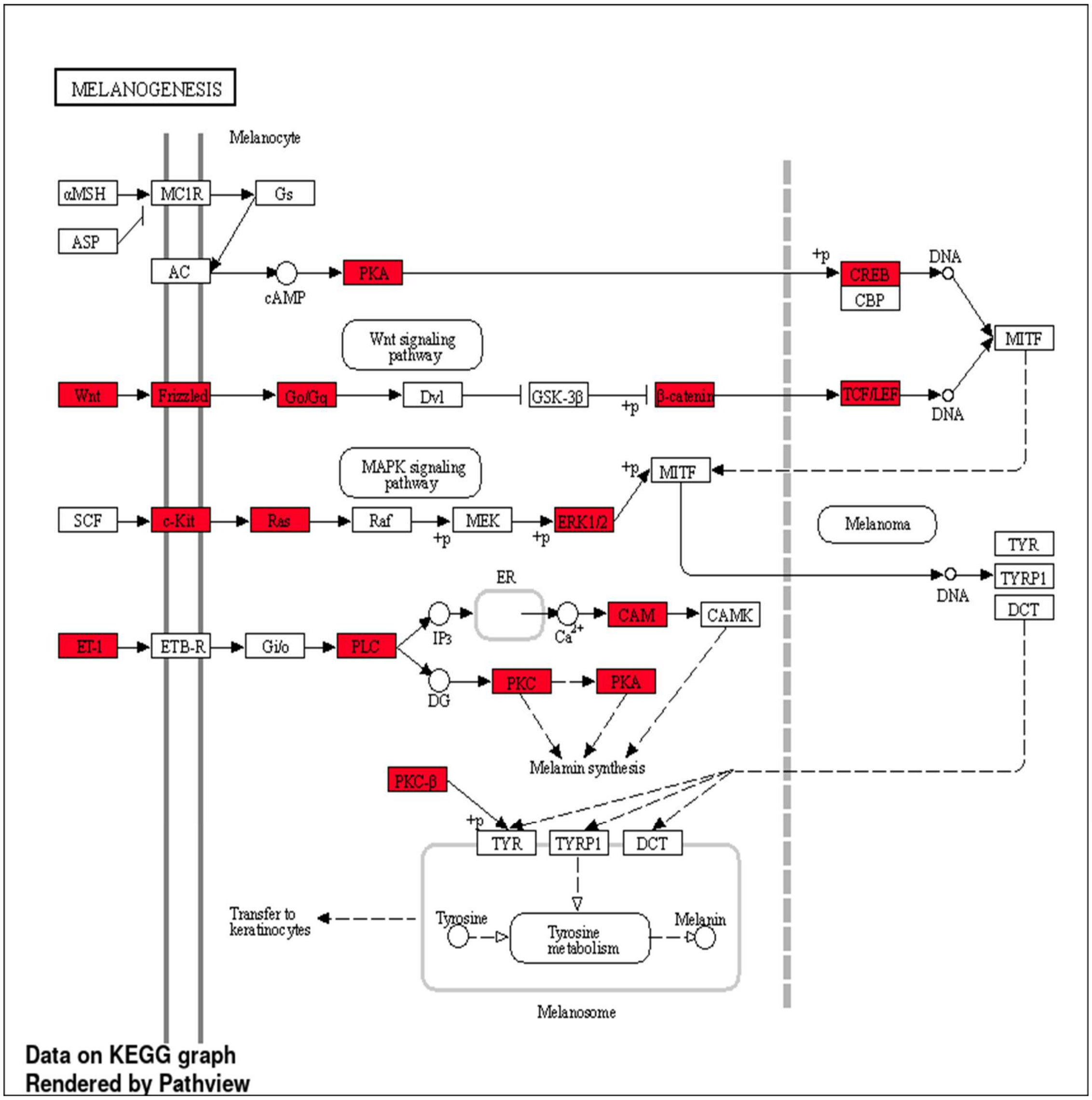


**Fig S35:** A diagram depicting the Melanogenesis pathway sourced from the KEGG pathway database, illustrating the relationship between the target genes regulated by the gga-mir-6552 paralog.
